# High rates of plasmid cotransformation in *E. coli* overturn the clonality myth and reveal colony development

**DOI:** 10.1101/2021.03.19.434223

**Authors:** Delia Tomoiaga, Jaclyn Bubnell, Liam Herndon, Paul Feinstein

**Affiliations:** Department of Biological Sciences, Hunter College, City University of New York, New York, New York 10065, USA; Manhattan Hunter College Science High School, New York, NY, USA; The Graduate Center Biochemistry, Biology and CUNY-Neuroscience-Collaborative Programs, City University of New York, New York, New York 10065

## Abstract

The concept of DNA transfer between bacteria was put forth by Griffith in 1928. During the dawn of molecular cloning of DNA in the 1980s, Hanahan described how the transformation of DNA plasmids into bacteria would allow for cloning of DNA fragments. Through this foundational work, it is widely taught that a typical transformation produces clonal bacterial colonies. Using low concentrations of several plasmids that encode different fluorescent proteins, under the same selective antibiotic, we show that *E. coli* bacteria readily accept multiple plasmids, resulting in widespread aclonality and reveal a complex pattern of colony development. Cotransformation of plasmids occurs by either CaCl_2_ or by electroporation methods. A bacterium rod transformed with three plasmids - each expressing a high level of a unique fluorescent protein - and replated on agar, appears to reassign a random number of the three fluorescent plasmids to its daughter cell during cell division. The potential to simultaneously follow multiple lineages of clonally related bacteria in a bacteria colony would allow for mosaic analysis of gene function. We show that clonally related bacterium rods self-organize in a fractal growth pattern and can remain linked during colony development revealing a potential target against microbiota growth.

## Introduction

Manipulation of *E. coli* colony development by DNA transformation has not been studied. *E.coli* colonies are generally viewed as a collection of bacterium with non-cell autonomous signaling rather than a developing organism. Mutant bacterial rods are studied for their ability to form a colony or biofilm. Typical competition studies are performed by the juxtaposition of large numbers of wild type and mutant bacterial rods. The ability of making mutants mosaically distributed in a developing colony has not been exploited. Here we have taken a *de novo* analysis of DNA transformation and uncovered that single bacterial rods can readily take up multiple plasmids and reveal for the first time how *E. coli* colonies develop. We show the potential to exploit this process for the study of mutations mosaically distributed within the development of a wild type colony.

Transformation of *E. coli* with DNA plasmids is a step for amplifying DNA fragments of interest, during molecular cloning. A typical procedure for generating DNA plasmids entails the cutting of a parent “vector” backbone carrying resistance and an origin of replication with a restriction enzyme along with alkaline phosphatase treatment to prevent vector re-closure. This vector is subsequently ligated with a fragment of interest. Other methods to incorporate an insert of interest also exist, such as ligations of PCR products with **A** overhangs directly into vectors that contain T overhangs (Promega) and DNA pairing (Gibson, TOPO); Ligated DNA is then transformed into “competent” bacteria, cells that have been pretreated to readily take up DNA.

Typically, DNA transformation is a two-step process: 1) pretreatment of bacteria with CaCl_2_ (or variations thereof) or with water and 2) subsequent heat shock or electroporation, respectively. Pretreatment of bacteria makes them highly competent^1^. Transformation efficiency is measured in transformants or colony forming unit (cfu) per μg DNA used, this being calculated at 10^9^ cfu/μg for CaCl_2_ pretreatment and subsequent heat-shock. The competency efficiency is about 10^11^ cfu/μg for the water pretreatment/electroporation method. Competency status of bacteria is usually established with 1 to 30 nanograms (ng) of supercoiled DNA, which is comparable to the quantity used for DNA cloning procedures. One nanogram of a typical supercoiled 4 kb plasmid contains about 2.3 x 10^8^ molecules and would yield a maximum of 10^6^ colonies. Hanahan demonstrated that transforming more than 35 ng of DNA yielded similar numbers of colonies as 1 ng, suggesting that the number of plasmid molecules is at threshold relative to the number of competent bacteria ^1^. Finally, the hallmark of DNA cloning is that the methodology yields bacterial colonies from single bacterium that are said to be clonal. By, contrast co-transformation of two different plasmids with two selectable markers is possible, but does not lead to development of mosaic colonies.

### Do standard methods in DNA cloning lead to clonal colonies?

In the past, our laboratory has generated large plasmids of up to 20 kb by standard DNA cloning methods to create targeting vectors for homologous recombination in embryonic stem cells ^2, 3^. Targeting vectors (TV) DNA consisted of a vector backbone and two arms of genomic DNA (∼4 kb each) with a PacI site between the arms. We incorporated reporter DNA cassettes at the PacI site by digestion of a second plasmid and gel isolation of a ∼5 kb IRES-tauLacZ (ITL) reporter fragment from its 3 kb vector backbone. Ligation of ITL PacI insert into a PacI digested TV yielded dark blue colonies due to the LacZ gene in the cassette after addition of X-gal. Such colonies were subsequently grown in liquid culture followed by DNA isolation, sequencing and restriction analysis.

We observed that *E. coli* cultures growing a correct clone reached saturation very slowly (up to 24 hrs) and produced low quantities of plasmid DNA. However, many uniformly dark blue colony cultures became confluent much faster (12 hrs) and upon restriction analysis and DNA sequencing were revealed to be a religated PacI vector without ITL, which did not produce dark blue colonies upon retransformation. We found this result especially surprising as the 5 kb (ITL) and 3 kb backbone vector fragments were easily separable by gel electrophoresis. Thus, any contamination of the ITL insert with its 3 kb backbone vector would be very low. This persistent anomaly forced us to grow multiple clones and throw away the rapidly growing cultures, which led to 100% identification of TVs ligated to ITL insert.

Our observation could be explained in several ways: 1) our isolated bacterial colonies with TV+ITL sat next to single bacterium transformed with the 3 kb vector; thus the resulting colony that appeared pure was a mix of two clones, or 2) the initial bacterium transformation event occurred with both TV+ITL and with a recircularized 3 kb backbone vector that once carried ITL, where the TV+ITL plasmid appeared to be the predominant plasmid. However, upon growth in liquid culture the recircularized plasmid of the 3 kb vector outcompeted the slower growing TV+ITL plasmid. Therefore, in the first instance it would be an issue of trivial contamination while in the second case it would be a contamination problem by non-clonality, especially disturbing since contamination occurred from a cotransformation with a second plasmid in very low abundance.

### A direct test for clonality

To distinguish between these two possibilities we constructed a set of nearly identical plasmids differing only in their expression of several fluorescent proteins (mCerulean, Venus - both derivatives of *Aequorea victoriae -* and mCherry, aka Cherry, from *Discosoma* sp.) that can easily be distinguished by confocal microscopy ^4^. To generate plasmids, we subcloned into pGEM-T easy (ColE1 origin) PCR products containing a constitutively active lactamase promoter (Lam→) upstream of the start codon of a fluorescent proteins (XFP). Each of these plasmids also contain a lactamase resistance gene (*lam*) in its backbone (see Data File S1).

We initially performed a straightforward CaCl_2_ transformation experiment by either combining all three plasmids post-heat shock (Separate Transformation) or prior to heat shock (Mixed Transformation). We expected to see the same results from both experiments: circular colonies, each expressing only one fluorescent protein. But, this pilot experiment led to two unexpected observations. From the Separate Transformation, we found that most colonies revealed Cerulean or Venus or Cherry fluorescence alone (Figure 1A-E), however, some colonies contained multiple fluorescent bacteria and maintained the typical circular shape (Figure 2A-D). In some circumstances non-circular colonies were observed that were clearly derived from distinct populations of transformants (Figure 2E-H); We observed torturous routes taken by one fluorescent population within another one (Figure S1A, S1C-F); when these resuspended colonies were diluted serially and aliquots of those dilutions were plated in order to obtain plates with single colonies that grew up each from a single cell, distinct fluorescent colonies emerged. An additional colony type was easily observed in the Mixed Transformation, appearing to contain different combinations of fluorescent proteins at different fluorescent intensities in patterns that did not resemble sectors or concentric rings (Figure 1F-J, 2I-L, 2L1a-d, 2L2a-d). This pattern is different than previously described ^5–7^. When we dispersed and replated these multiple fluorescent colonies at limiting dilutions, the fluorescent mixtures were maintained in nearly all-resulting colonies suggesting that three plasmids were common to each bacterium (Figure S1B, S1G-J). Based on this result, it was difficult to argue against the idea that a triple transformation event had occurred.

**Fig. 1.**
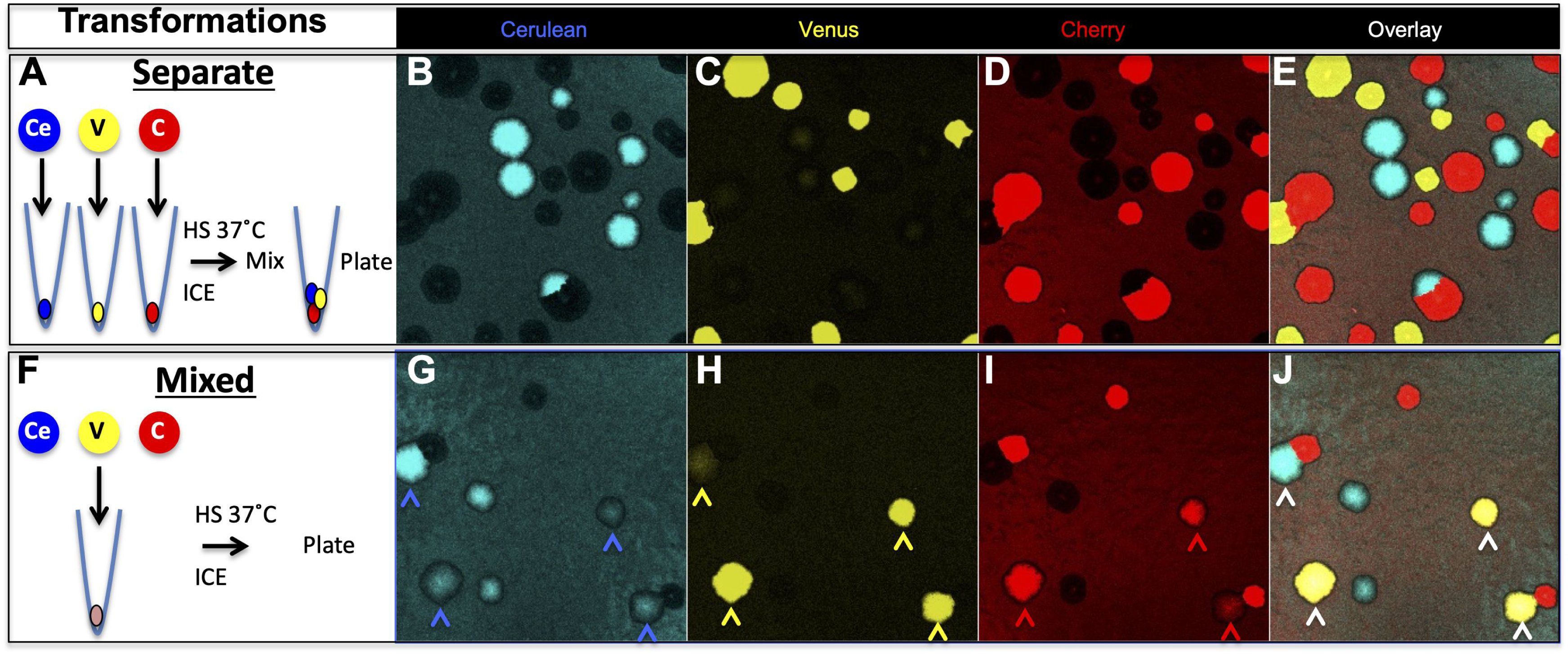
Transformation with three plasmids separately or in mixed. **A.** Three lactamase promoter (Lam→) constructs each expressing a different fluorescent protein (Ce-Cerulean, V-Venus or C-mCherry) were transformed separately into DH5 (*E.coli*. **B, C, and D.** Colonies observed after 16 hrs of growth (all plasmids have the same resistance). Overlay in **E** shows no colony has double fluorescence using saturating laser excitation. **F.** Three constructs cotransformed simultaneously. **G, H, and I.** Colonies observed after 16 hrs of growth (all plasmids have the same resistance). **J**. Four of the nine colonies in this visual field show coexpression (white arrows) using saturating laser excitation. Three of these colonies express all three fluorescent proteins and one only Cerulean and Venus (see colored arrows in **G**, **H**, **I**).

**Fig. 2.**
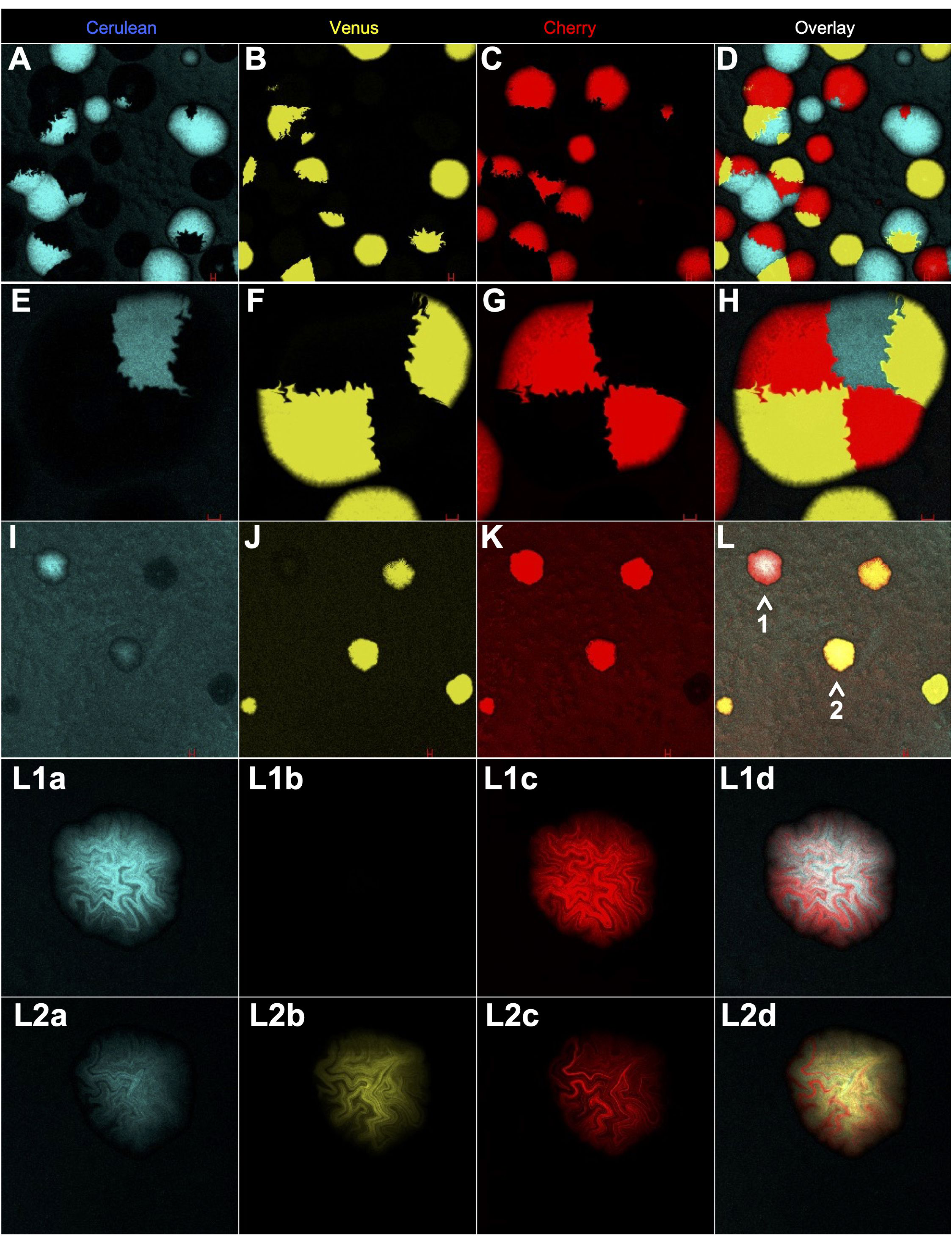
High density plating produces mixed segregating colonies whereas coexpressing colonies are mosaic. **A-C.** Low magnification image of many segregated round colonies. **D** is overlay. **E-G,** High magnification image; one colony made up of five separate transformants using saturating laser excitation. **H** is overlay. **I-K.** Low magnification image of mixed transformations; four of five colonies coexpress using saturating laser excitation. **L** is overlay (arrows 1 and 2 are two coexpressing colonies). **L1a-c** and **L2a-c** are high magnification image of colonies 1 and 2, respectively in L observed with non-saturating laser excitation. **L1d and L2d** are overlays clearly revealing reciprocal mosaicism of fluorescence.

### Direct evidence for cotransformation

Validation of cotransformation by genetic complementation would firmly establish the cotransformation phenomena. Recent advances have shown GFP protein can be split into two portions capable of re-annealing: beta strands 1-10 and 11 only ^8^. In such a circumstance, robust fluorescence can only be observed when both portions of the GFP are expressed in the same cell. Thus, we created expression vectors that express each portion separately using sfGFP ^9, 10^ (see Data File S1) and determined whether green fluorescence could be observed. Such fluorescence would parallel patterns of *E.coli* development observed by the three plasmid cotransformation experiments (Figures 2L1 and 2L2), since the only fluorescence that could be observed will occur from stable expression of both plasmids in the same bacteria. In fact, this is precisely what we observed. No-fluorescence was seen with either portion of sfGFP alone (Figures 3A and 3B), but when both were coexpressed, fluorescence was readily detected (Figure 3C) and in all bacterial colonies had fluorescence in a mosaic pattern (Figure 3D). No colony could be observed that was uniformly fluorescent, which also suggests that plasmid equilibrium is a robust competitive process during the earliest rounds of bacterial divisions. By contrast, robust fluorescence (Figure 3C and 3D) was not observable in colonies there were a mixture of sfGFP1-10 and GFP11 bacteria, showing that bridges between colonies cannot reconstitute full fluorescent sfGFP molecule.

**Fig. 3.**
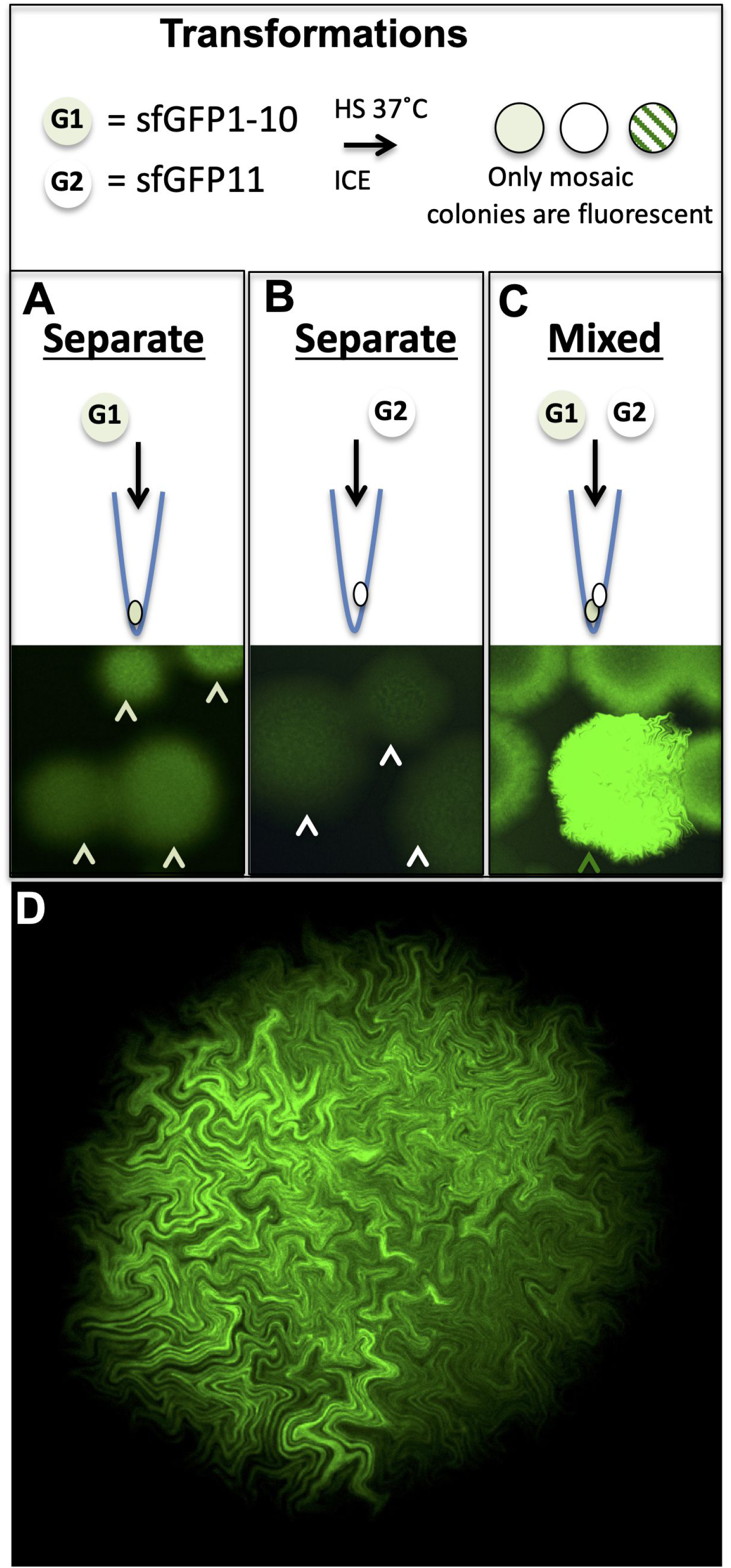
Contransformation complementation of split sfGFP in *E.coli*. **A-C.** Saturating laser excitation of *E.coli* colonies transformed with Split sfGFP expression plasmids. **A.** sfGFP 1-10 (G1), incomplete beta barrel structure yields faint fluorescent colonies **B.** sfGFP 11^th^ beta strand (G2) no fluorescence. **C.** G1+G2 reveals high levels of expression. **D.** Under non-saturation imaging all G1+G2 fluorescent colonies reveal only mosaic expression. This could only be achieved by an initial transformation event having taken up both plasmids, followed by stochastic loss/reduction of either G1 or G2 plasmids.

### At least 11 plasmids can be simultaneously cotransformed

In order to determine cotransformation events, we set up a PCR screen for each fluorescent plasmid. Venus and Cerulean coding sequences are nearly identical, so we altered the codon usage for Venus at its most divergent point to Cerulean to generate specific oligomers and we confirmed by PCR the presence of all three plasmids. Because we readily observed cotransformation of three different plasmids in a bacterial colony, we surmised that it would be possible for even more plasmids to cotransform.

We constructed a set of unique Venus expression plasmids by the addition of nucleotide sequence specific tags after the stop codon to extend our analysis (see Data File S1). In addition, we switched to the brighter blue fluorescent protein mTeal, aka Teal, from *Clavularia* sp. Here, we made a mixture of 10 Venus plasmids (V1-V10), 1 ng each, along with 1 ng of our Cherry plasmid and cotransformed the 11 ng mixture. However, prior to plating, we added 1 ng of Teal expressing plasmid to control for stickiness of a plasmid that might confound our PCR test (Figure 4A). It is important to note that Teal, Venus, and Cherry are from three different organisms and share minimal nucleotide homologies.

**Fig. 4.**
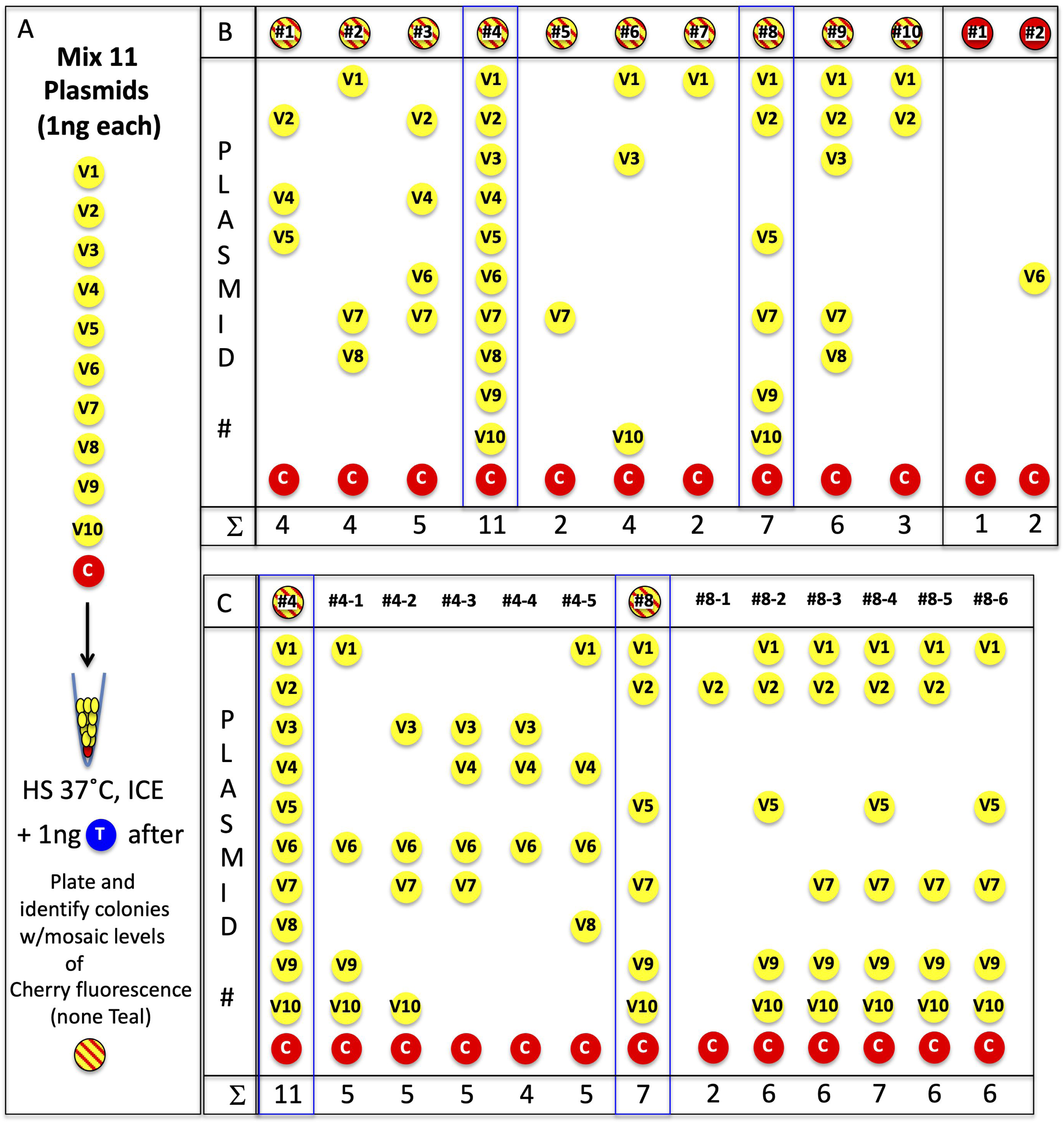
*E. coli* can take up at least 11 plasmids. **A.** 10 versions of the Lam→Venus plasmids were created, each separable by PCR. 1 ng of each were mixed with 1 ng of Lam→Cherry and cotransformed. Following cotransformation, 1 ng of Lam→Teal was added prior to plating (all 12 plasmids carry same resistance). **B.** PCR analysis of 10 minimally Cherry fluorescent mosaic colonies and two colonies that appeared Cherry fluorescent only. Lam→Cherry was detectable by PCR, whereas Lam→Teal was not detectable. Colony #4 was positive by PCR for all 11 PCR primer sets and #8 was for 7 PCR primer sets. **C.** Colonies #4 and #8 were subcloned and a few colonies were tested by PCR. An average of 5 plasmids were stably maintained.

Venus fluorescent colonies were easily identified with Cherry fluorescent mosaicism by microscopic inspection; no Teal fluorescence was observed in any colony. PCR analysis of 10 mosaic colonies revealed the presence of many combinations of Venus plasmids along with the Cherry plasmid, averaging 5 unique plasmids per bacterial colony (Data File S1); No Teal plasmid could be identified as anticipated (Figure 4B). We were surprised that one colony (#4) contained all 11 plasmids and another (#8) contained 7 plasmids. Both of these colonies were replated at limiting dilutions to obtain sparse single colonies. These subclones reanalyzed by PCR: Colony #4 subclones, #4-1 to #4-5, revealed differing subsets of the 11 Venus plasmids and none contained more than 5 plasmids. However, colony #8 subclones, #8-1 to #8-6, revealed that 5 out of 6 colonies contained at least 6 plasmids with #8-4 having all of the original 7 plasmids, showing the surprising stability of these multiple Venus plasmids (Figure 4C). Thus, at least 11 distinct plasmids could be cotransformed into a single bacterium using 11 ng of plasmid DNA. It should be noted that 11 ng of plasmid DNA is a significant lower number than most laboratories use in transformation experiments. Based on the ease of this observation, it is likely that many more plasmids can be cotransformed.

### A smaller plasmid outcompetes a larger one

The above observations suggested that our hypothesis of cotransformation of TV+ITL and the recircularized donor vector must be true. To directly confirm our original observations we took two plasmids, one expressing Cherry (3,805bp) and a targeting vector containing the ITL cassette (∼17.5 kb). We mixed 1 ng of each together and introduced them into bacteria using the CaCl_2_ transformation method. We obtained 151 colonies: 146 Cherry fluorescent positive by microscopy and 5 that were devoid of Cherry fluorescence. As expected all 5 of the non-Cherry fluorescing colonies amplified early in the Sybr Green based qPCR detection process solely with primers specific for ITL. However, 4 out of 146 Cherry fluorescing colonies also amplified with primers specific for ITL, albeit late in the Sybr Green based qPCR detection process, suggesting they were in much lower abundance (Data File S1). Not surprisingly, replating and repeating the PCR on subclones from the 4 colonies failed to detect any residual ITL. Thus, we now confirm our hypothesis and show directly that the fast replicating 4 kb Cherry plasmid outperformed the larger plasmids. We show that 2 ng total plasmid DNA (1.25 ng molar equivalents of replicons: 1ng of the Cherry plasmid and 0.25 ng of a ∼4 kb plasmid DNA for the 1 ng of ∼17.5 kb plasmid) can result in observable cotransformation rates of 2.6% (4 out of 151 of the total bacterial colonies). This rate of cotransformation to be more common than had been described in the literature.

### CaCl_2_ mediated cotransformation of plasmids is a common occurrence

PCR is a sensitive way to detect cotransformation, however, two confounding variables make it difficult to use as a quantitative test of absolute cotransformation rates. First, it is not possible to perform PCR on an entire colony, thereby every cell in a colony cannot be reliably analyzed. Second, even if all bacteria could be assessed, PCR cannot differentiate between a mosaic bacterium carrying multiple plasmids or two neighboring bacteria each carrying different plasmids. We had initially confirmed our findings of cotransformations with multiple plasmid DNAs by a strategy of limited dilution and replating and subsequent reanalysis by confocal microscopy. In general, PCR identification of multiple plasmids from our transformations with low DNA concentrations, resulted from cotransformation events rather than contamination from a neighboring colony.

Unfortunately, our Lam→XFP plasmids did not fluoresce with enough intensity to provide us with single bacterium resolution and therefore we could not assess the fates of any single bacterium hiding within a bacterial colony. However, while shuttling fluorescent DNA inserts during pGEM-T cloning we observed a single strong Venus expressing bacterial colony, about 6-8x more fluorescent intensity (Hi→Venus) during colony formation than our original Lam→Venus version. This new plasmid, to be described in detail elsewhere, relied on the lac operon promoter to create constitutively expressed fluorescent proteins throughout colony formation. Using this vector template, we substituted Venus with other XFPs to generate a series of Hi→XFP plasmids (*lam*): Hi→Cerulean, Hi→Teal, Hi→Venus (codon altered) and Hi→Cherry. We analyzed single bacterium fluorescence for all the four XFPs and only Hi→Cerulean fluorescence was not reproducibly observed in single bacteria. Hi→XFP mixtures (Teal/Venus/Cherry) provided the foundation for our endeavor to understand cotransformation rates where single bacterium resolution can be readily imaged (Figure 5B, S5A-D, Movie S3).

**Fig. 5.**
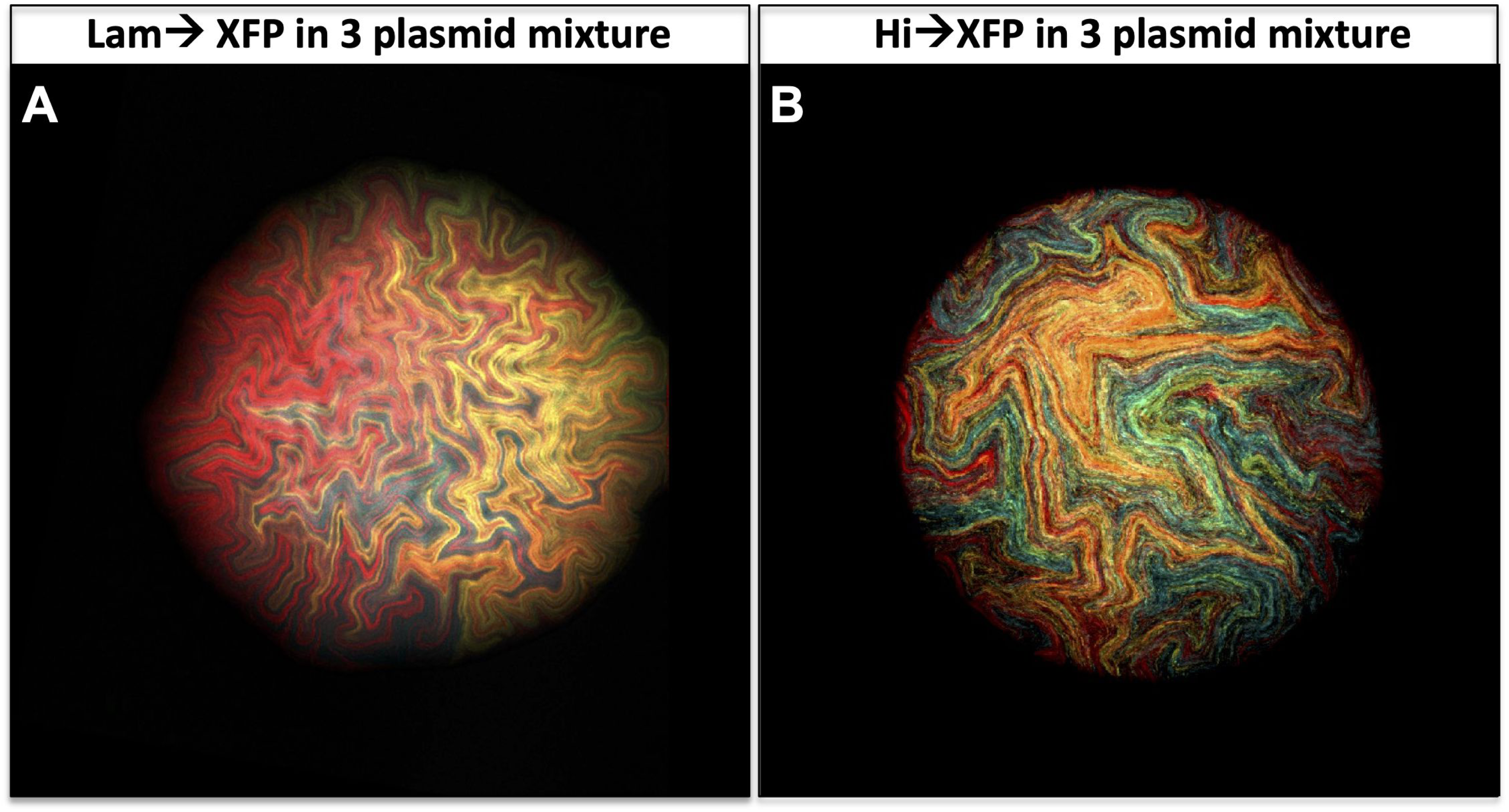
High expression fluorescent plasmids allow for single bacterium fluorescent detection. **A.** A triple fluorescent colony (Lam→Teal, Lam→Venus, and Lam→Cherry) from a projected Z-Stack reveals “rivers” of fluorescence. In contrast, **B.** A triple fluorescent colony (Hi→Teal, Hi→Venus, and Hi→Cherry) from a projected Z-Stack, reveals fluorescence rod shapes throughout the image. These rods are indicative of single bacterium as we observe in Fig.7. Typical colonies after 24 hrs of growth are 500µm in width.

The first version of our cotransforming assay was perfomred by mixing 0.1 ng of Hi→Venus and 0.1 ng of Hi→Cherry plasmids followed by the addition of 1 ng of Hi→Teal plasmid post heat shock. PCR analysis on 95 colonies and subsequent replating by limited dilutions of two candidate cotransformed bacterial colonies revealed 1 out of 95 colonies was truly a double positive and 1 out of 95 was a mosaic colony of two separate transformants. No Hi→Teal could be amplified or visualized in the colonies. Thus, 0.2 ng total DNA yielded a ∼1% observed double positive (ODP) rate of cotransformation (Data File S1). However, we do not score for two Hi→Venus or two Hi→Cherry plasmids cotransforming (unobservable double positive, UDP) with the same plasmid. Taking into account these unobserved double positives (Data File S2), the estimated double positive (EDP) rate for 0.2 ng is higher and closer to 2% (Table 1). Our finding that only about 98% of colonies are clonal colonies when using 0.2 ng of plasmid DNA is crucial and dispels the current belief that CaCl_2_ transformation into bacteria yields clonal colonies (Table S1).

**Table 1.** Summary of cotransformation rates with multiple plasmids under different parameters that include the unobservable events. *The Kan→Carb column reflects concentrations for Carb only. In addition to this concentration, 1 ng of Kan plasmid was added during cotransformation experiments.

### Three plasmid transformations reveal high rates of aclonal colonies

A slight 0.1 ng increase in plasmid DNA to 0.3 ng (0.1 ng of each: Hi→Teal, Hi→Venus, and Hi→Cherry) yielded 3 out of 95 colonies with multiple plasmids or a 3% ODP rate (97% clonal). Again, we cannot score for double positive colonies of two Hi→Teal, Hi→Venus, or Hi→Cherry plasmids, so these rates are likely conservative. When calculating the unobserved events for three plasmid using 0.3 ng, the clonality rate dips to 94% with an EDP rate of cotransformations of 6% and no observed triple positives (OTP). Based on our ability to directly visualize single bacterium fluorescence and discern the difference between two colonies growing together and a true double positive bacterium, we finalized a high-throughput scoring method where 96 random colonies were picked into a microtiter dish, grown overnight and replated as 1μl droplets on a fresh plate and grown for a second night (Figure S2 and S3). This replating strategy accurately distinguished neighboring (mosaic) from double or triple positive colonies (Data File S1). We next tested 10 fold more plasmid DNA, 3 ng (1 ng for each of Hi→Teal, Hi→Venus, and Hi→Cherry), which yielded 9 double positives and one triple positive and increases the ODP cotransformation rate to 10% and the observed triple transformation rate (OTP) to 1%. Once again, the non-observable triple cotransformation events such as 3 plasmids of Hi→Teal, Hi→Venus or Hi→Cherry or paired combinations of one plus a different plasmid are not counted. The estimated rate of clonality drops to 82%, which is surprising and notable for the practicality of molecular cloning methods, as most laboratory transformation protocols are performed with more than 3 ng of DNA.

Finally, we tested transformation rates in the range used in standard cloning and retransformation, 30 ng of plasmid DNA (10 ng of each Hi→Teal, Hi→Venus, and Hi→Cherry plasmids). Our rates of clonality further decreased to 72% based on 20% ODP and 8% OTP colonies, but the estimated clonality was only 44%. Thus, 30 ng of plasmid DNA transformations using CaCl_2_ mediated transformation and plated on carbenicillin (Carb) selection plates yields low rates of clonal colonies, scored by microscopy. We repeated these observations over the course of months and at various concentrations and readily established a colony clonality rate relative to the amount of plasmid used (Table S1 and Data File S1). These observed clonality rates for the transformations were further reduced when we accounted for the unobservable events (EDP and ETP-Data File S2 and Table 1). Finally, performing transformations with 0.1 ng of 0.033 ng for each Hi→XFP using purchased, highly competent (10^9^ cfu/μg) bacteria, we found 3.3% ODP and 0% OTP colonies, similar to our 3% ODP and 0% OTP using 0.3 ng and lower than our 4% ODP and 1% OTP using 1 ng in our laboratory produced competent (10^7^ cfu/μg) bacteria. Thus, using cells 100 times more competent does not increase cotransformation rates by 100 fold (Data File S1) leading to the conclusion that cotransformation rates are dependent on the DNA concentration and not the competency status of the bacterial aliquot.

### Cotransformation frequencies also occur using kanamycin selection

Our analysis suggested that using standard cloning methods with calf intestinal phosphatase (CIP) treatment would reduce cotransformation rates that would otherwise be rampant with recircularized vector plasmids. We also observed mixed fluorescent colonies after transforming when performing simultaneous triple insert ligations (Cerulean, Venus and Cherry) into a CIP treated vector; These mixed clones did not contain vectors with multiple inserts. In all experiments so-far described, the plasmids that were cotransformed contained the same selection marker (*lam*).

The high rates of cotransformation suggested that the occurrence of mixed fluorescent colonies is solely dependent on DNA concentration and not plasmid selection by a resistance marker. To test this hypothesis in CaCl_2_ cotransformation experiments, we mixed plasmids with two different resistance markers, kanamycin (Kan) and carbenicillin (Carb)- *kan* and *lam*, respectively (Data File S1). Here we selected only for the Kan plasmid and asked if we could observe fluorescence from the Carb containing plasmid that might have be cotransformed. Initially, we cotransformed 1 ng of a ∼5kb plasmid containing a weak cherry (Lo→Cherry) fluorescent reporter (*kan*) with 2 ng of a Hi→Teal (*lam*) or 2 ng of a Hi→Venus ∼4kb plasmid (*lam*), 3 ng total. We randomly selected 96 colonies that grew on Kan only plates. Colonies were picked and subsequently grown in Carbenicilin media, which confirmed that Carb containing plasmids did indeed co-transform with the Kan containing plasmids. (Figure S4A-P). We observed 19% of the colonies were cotransformed (17 out of 96 Kan resistant colonies were highly fluorescent for Teal, without this plasmid being selected, and in a separate experiment 20 out of 96 Kan resistant colonies and were highly fluorescent for Venus, without this plasmid being selected, and both sets were weakly mosaic for Lo→Cherry). The clonality rate of 81% for 3 ng of total plasmid is similar to the 91% observed Carb resistance alone for 3 ng of plasmid DNA. Transformation rates were based on growth in Carb media followed by direct visualization.

We were in a position to ask if the ODP rates were a property of the total DNA transformed or simply on the Carb resistant plasmids. Cotransformation of 0.2 ng of Hi→XFP mix (*lam*) along with 1 ng of the Kan plasmid yielded 5% double positive colonies whereas 10 ng of the Hi→XFP mix (*lam*) yielded 40% double positives, 5% triple positives and 1% quadruple positives (Table 1; Figure S4Q-T and S5E-F). Because we cannot ascertain multiple Kan resistant colonies: double, triple, or quadruple positive colonies, estimated clonality for the total 1.2 ng and 11 ng cotransformations must be lower than 95% and 54%, respectively. Next, we cotransformed 11 ng of DNA, but with 1 ng of the Hi→XFP mix (*lam*) and 10 ng of the *kan* plasmid, which gave a 88% clonal rate, which is between the 0.2 ng and the 2 ng rates for the Hi→XFP and distant from 10 ng Hi→XFP rates (Table 1). Thus, cotransformation rates in this context were not dependent on the total amount of Kan resistant DNA used. It was surprising that selection on Kan plates did not affect rates of cotransformation of a Carb resistant plasmid. Finally, we created Hi→XFP-Kan ∼5 kb plasmids, by cloning a 1 kb *kan* resistant gene within the *lam* gene and tested if clonality rates under Kan selection differed from Carb selection. The main difference here is that these transformations are required to grow in nonselective media for at least 30 minutes prior to plating. To our surprise, combined ODP and OTP rates of cotransformation appeared quite low and plateaued at 13% for both 10 ng and 30 ng Hi→XFP-Kan mixes (Table 1). It is possible that these cotransformation rates were lower than the Carb plasmids due to their 25% increase in plasmid size.

### Cotransformation frequencies by electroporation are temperature sensitive

Thus far we have focused on the standard Hanahan CaCl_2_ transformation technology and hypothesized that cotransformation may be tightly associated with this poorly understood and unknown mechanism. By contrast, electrocompetent bacteria take up plasmids through a water environment during the process of electroporation (EP)^11–13^ and may be immune to cotransformation. Using our Hi→XFP Carb resistant plasmids, we show ODP rates were fairly low for 1 ng to 30 ng DNA concentrations, ranging between 3-6% aclonality rates. In addition, OTP rates were almost non-existent, 1 out of 1042 colonies analyzed. We were surprised by these results and decided to electroporate bacteria with higher concentrations: 300 ng mixture of Hi→XFP, achieved 36% ODP and 10% OTP rates, driving the clonality down to 54% of the colonies. Thus, the electroporation process is not resistant to high cotransformation rates.

Electrocompetent bacteria are normally prepared by several washes in ice-cold water, followed by maintaining bacteria on ice in water prior to EP in cuvettes kept on ice. These chilled cuvettes are at an elevated temperature relative to the chilled bacteria in a liquid environment. Thus, adding the bacteria/DNA mixture prior to EP has the potential to alter the bacteria making them less competent or able to take up DNA by EP. We asked if maintaining the bacteria in a colder state by keeping cold bacteria in *freezer*-chilled cuvettes would alter cotransformation rates. Indeed, transformation rates rose dramatically with the identification of triple transformed colonies in the 1 ng Hi→XFP mix (2 out of 168 colonies) and in the 10 ng Hi→XFP mix (5 out of 204 colonies) as compared to a single observation among 1042 colonies from the chilled cuvettes. Thus, if bacteria are maintained in a cold environment during the electroporation, the clonality rates of Hi→XFP plasmids of 97% for 1 ng and 83% for the 10 ng are similar to those observed during CaCl_2_ treatment (Table 1, S1 and Data File S1). Therefore, cold, but not ice-cold, cuvettes can limit cotransformation rates.

### CaCl_2_ transformation is dependent on plasmid association with the bacterium

Using our Kan selection experiments, we show that cotransformation rates are not dictated by the total amount of plasmid DNA in a transformation reaction. Because the mechanism of CaCl_2_ transformation is unknown, we employed our Hi→XFP plasmids to test if the plasmid DNA-CaCl_2_ precipitates were associated with the bacterial membrane during the process. If yes, could we wash off the DNA from the membrane of competent cells prior to heat shock with CaCl_2_, water or Tris-EDTA (TE)? Thus, we designed an experiment to reveal any residual Hi→XFP plasmids by creating two 30 ng transformation conditions: An incubation with a mix of 10 ng each Hi→Teal, Hi→Venus, and Hi→Cherry (Hi→XFP-Mix) followed by 2x ice-cold CaCl_2_ washes or three separate 10 ng incubations of Hi→Teal, Hi→Venus or Hi→Cherry (Hi→XFP-Sep) followed by 2x ice-cold CaCl_2_ washes and brought together into the same tube after washes. Cotransformation rates of 30 ng Hi→XFP-Mix colonies were at 27% despite 2x ice-cold CaCl_2_ washes, and 1% for the Hi→XFP-Sep colonies. Thus, the DNA appears to be tightly associated with the bacteria and cannot be washed off by additional CaCl_2_. We followed up on this experiment by using two washes with room temperature water (instead of CaCl_2_), which we expected would solubilize the potential plasmid-CaCl_2_ composites associated with an individual bacterium. However, transformation still occurred, but cotransformation rates for 30 ng Hi→XFP-Mix were lowered to 10% whereas the 30 ng Hi→XFP-Sep were reduced to 0%. But, no transformations occurred for the 2x room temperature TE washes. This latter result seemed inconsistent with the 2x RT water washes, which not only allowed for transformations, but also non-clonal colonies. So either the TE was solubilizing DNA-CaCl_2_ aggregates or maybe the RT water washes themselves might not be “washing” the DNA of the bacteria, but rather providing a heat shock to the bacteria prior to the placement at 42°C for 1 min; the 42°C (2^nd^ heat shock) might cause an increase in lethality due to the extended time that bacteria would be in the presence of elevated temperatures. Thus, we reduced the time at heat shock to just under a minute. CaCl_2_ transformations with 2x CaCl_2_ washes and reduced incubation at 42°C led to increased cotransformation rates up to 40% for the 30 ng Hi→XFP-Mix, and 3% for the 30 ng Hi→XFP-Sep colonies. In stark contrast to the room temperature water washes, the ice-cold water washes yielded no transformations and may have indeed washed off the plasmid from the bacterial membrane. We concluded that the plasmid DNA is loosely associated with CaCl_2_ treated bacterial membranes prior to heat shock, but unlikely to precipitate the cotransformation event as we observe cotransformation by electroporation without the presence of CaCl_2_ (Table S2 and Data File S1).

### Bacterial colony development suggests complex developmental processes

Finally, we sought to understand how our mixed colonies obtained their patterning. A large segment of the archetypical *E. coli* literature implies that bacteria colony formation occurs in subsequent expansions of concentric circles ^5–7^. However, our mixed colonies exhibited fractal patterns of development similar to early stages of replicating bacteria ^14–16^. To test if the fractal colony growth pattern is independent of constitutive high fluorescence expression, we decided to express our fluorescent proteins using an arabinose inducible plasmid system with the ColE1 origin of replication and to repeat the cotransformations, inducing XFP expression after colonies had formed. Not only were triple transformants and fractal patterns observed, but we also found that fluorescent intensity varied amongst a set of concentric rings within the bacterial colony, suggesting that differential distance from the colony center dictates gene expression patterns. (Figure 6A-H).

**Fig. 6.**
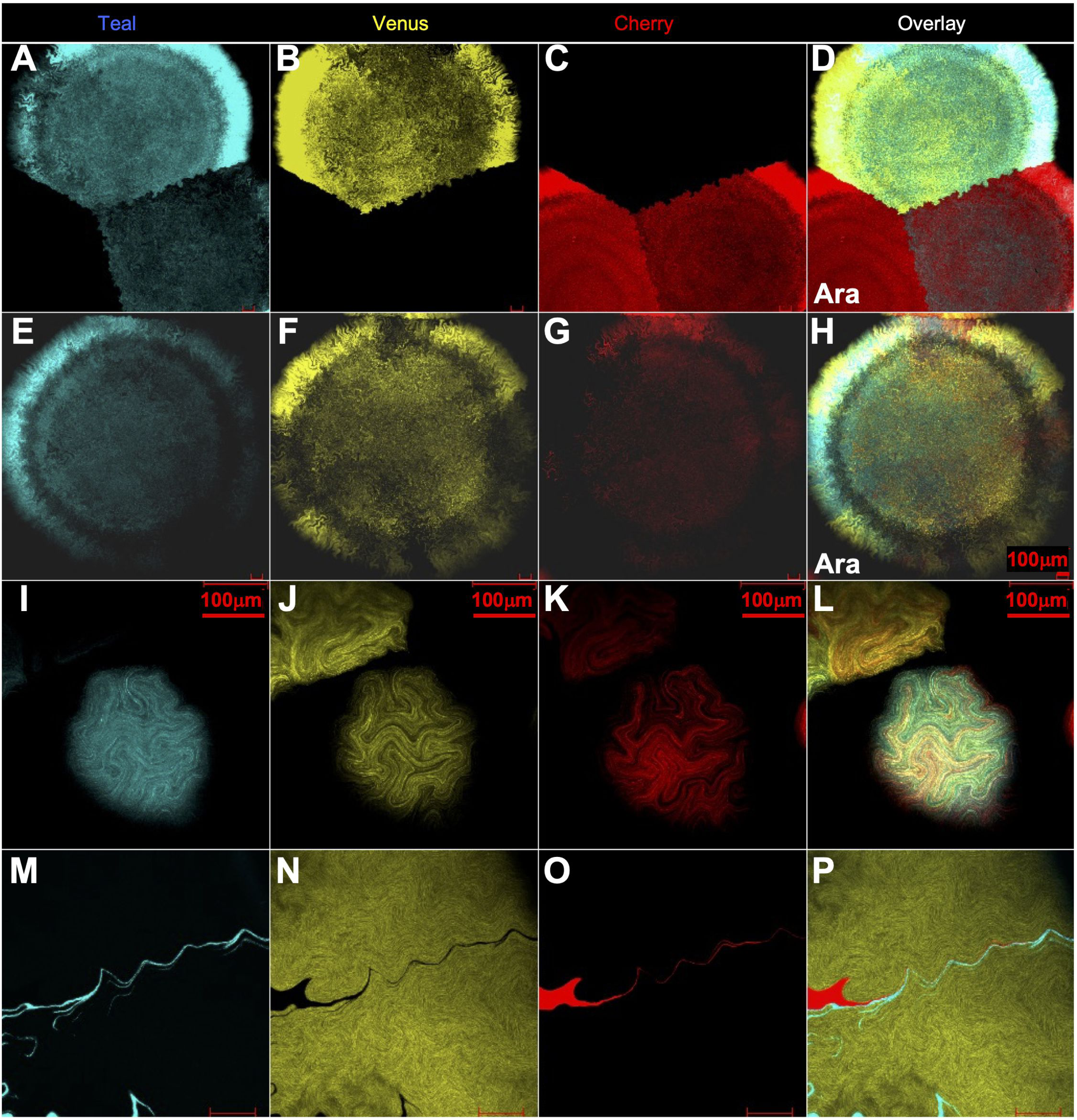
Mosaic fluorescence observed by induction or during colony development. **A-C,** L-Arabinose induced fluorescence from Ara→Teal and Ara→Venus coexpressing colony, Ara→Teal and Ara→Cherry coexpressing colony and Ara→Cherry colony. **D** is overlay. **E-G,** L-Arabinose induced triple fluorescence colony (Ara→Teal, Ara→Venus and Ara→Cherry). **H** is overlay. All colonies reveal waves of fluorescent expression as well as mosaic expression in coexpressing colonies. Bacterial colony is 2mm in diameter. **I-K** From triple fluorescent colony expressing Hi→Teal, Hi→Venus, and Hi→Cherry at <u>10hrs</u> of growth and 150µm in width. **L** is overlay. **M-O,** a Hi→Venus colony that contains bacteria for Hi→Teal and Hi→Cherry.

In an effort to observe how these Hi→XFP multifluorescent colonies emerge, we set up an extended time course to analyze how a single triple fluorescent bacterium forms a colony (Figure 7). Initially, this colony-founder bacterium divides in a fractal pattern and, after a few hours, different lineages of fluorescent bacteria emerge, presumably due to differential plasmid loads of the Hi→XFP plasmids (Figure 7; Figure S7 and S8; Movie S1A, S1B and S2). Colonies are not initially round, but after about 10 hours of the fractal growth pattern, the colonies achieve a more roundish shape; under high power magnification, the edges of the colonies are not in the shape of a perfect circle but rather look uneven like the gyri and sulci observed in cerebral cortex. Thus, the fractal growth pattern is still present when a colony is fully formed (Figure 6I-L; Figure S6).

**Fig. 7.**
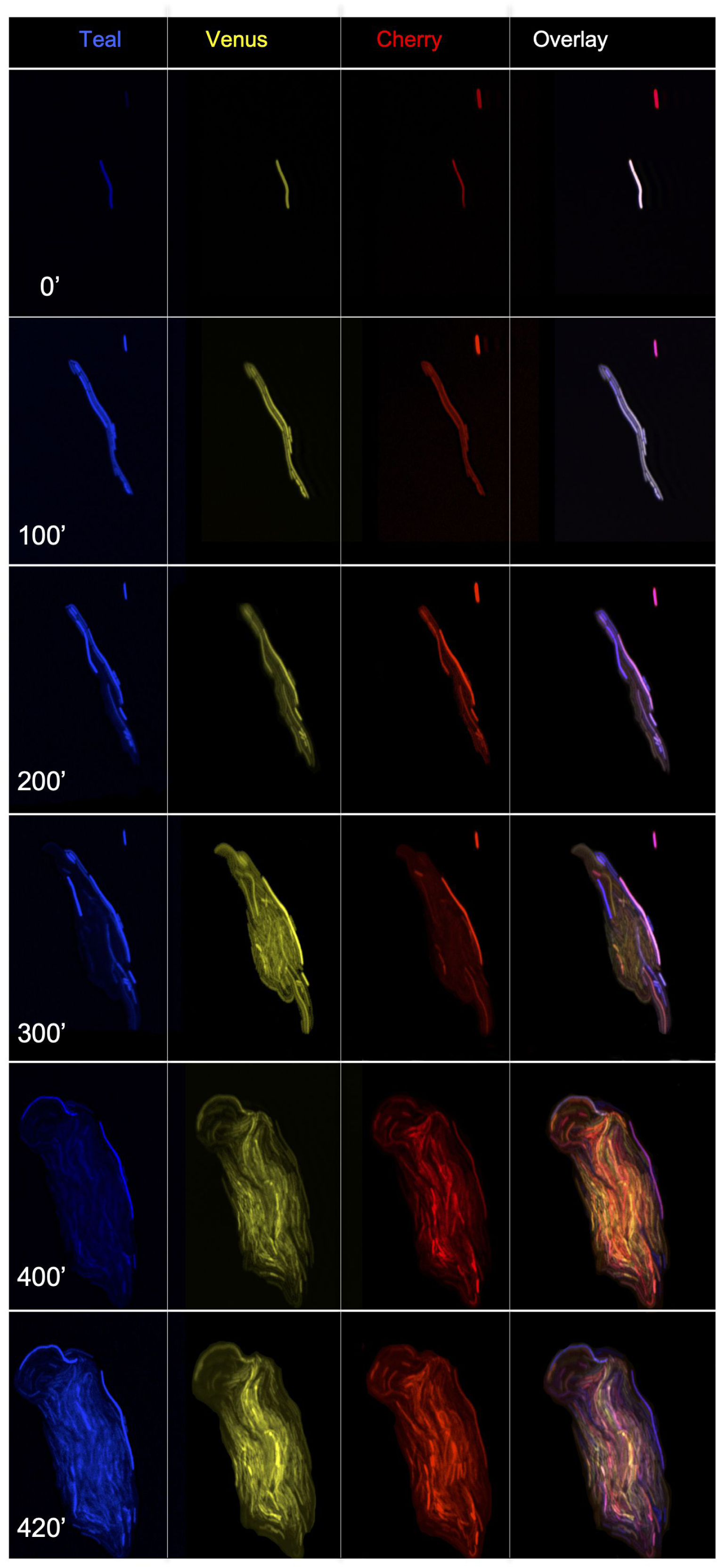
Time course of triple fluorescent colony development 0 minutes to 420 minutes. A triple fluorescent bacterium (A) expressing Hi→Teal, Hi→Venus, and Hi→Cherry was imaged every 20 minutes for 420 minutes. All 20 minute timepoints are in **Fig. S7** and **Movie S1A.** Mosaicism is quickly revealed after a few cell divisions.

While following the growth pattern of our *E.coli* bacteria, we arrived at two important findings. First, many of the colony-founder single bacteria failed to divide and were eventually swallowed up by a colony forming in close proximity. This can explain how mosaic bacterial colonies containing other bacteria units form. Second, we often observed the initial colony-founder bacterium producing replicates, but failing itself to divide without disappearing, creating a point of nucleation for the colony. Finally, our lineage tracing experiments revealed fluorescent bacterium that had divided and stayed attached to each other after replication (Figure 6M-P) creating a chain of linked progeny^17–21^. These linked chains could be observed in 3D (Movie S3 and S4). We concluded that our multi-fluorescent assay can be used for identifying genes involved in maintaining progeny linkage during replication, relevant to chemotaxis and lipid secretion behaviors of bacteria.

### Mosaicism to study development

In order to study mutant bacteria it is important for those mutations to allow for colony development. It is likely that large numbers of mutants could survive when surrounded by wild-type bacteria secreting or contributing non-autonomous signals. Our proposed mosaic colony screens will determine how colony development are affected when neighboring bacteria are mutated. In particular, mutations that disrupt robust fractal patterning would be most readily identified. There are several types of screens using cotransformation to place multiple plasmids in the same bacterium. We envision two types of mosaic analyses to study these event (Figure 8A and 8B): 1) A two population screen: where one plasmid reflects wild-type bacteria segregating from bacteria under or over-expressing a mutant protein. 2) A three-population screen: where each plasmid alone produces a phenotype and a third phenotype is observed when both plasmids are stably expressed in the same bacteria. It should be noted that all experiments described here are with the current plasmids, which constitutively express our proteins of interest. Other versions of these plasmids could yield distinct populations as well^22^. We foresee that adding the ability of protein induction to these schemes further allows bacterial colonies to prosper, until the mutation is imposed on its development.

**Fig. 8.**
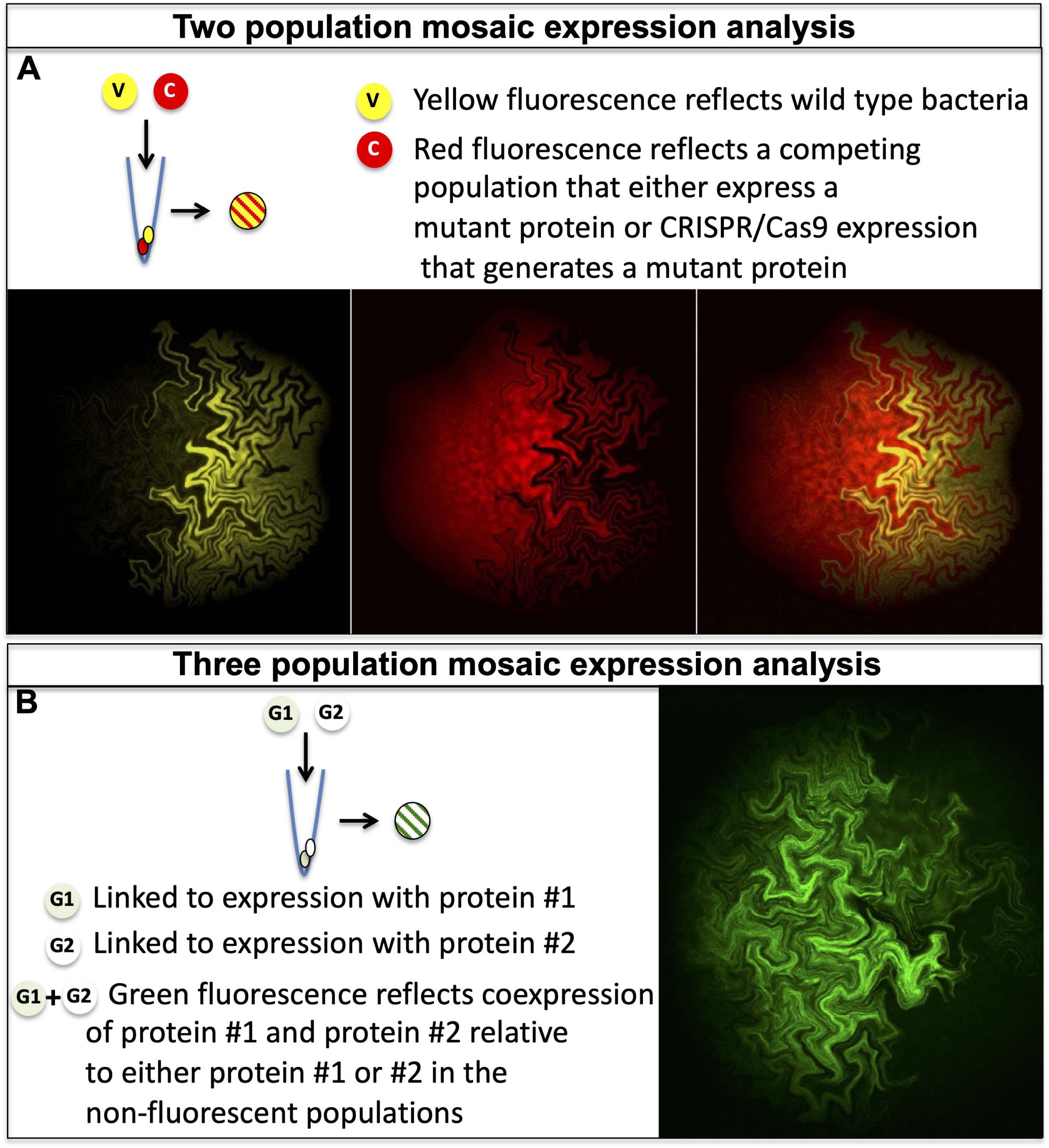
Proposed schemes for mosaic analysis of mutant *E.coli*. **A.** Two population mosaic expression analysis. In this genetic screen a plasmid expressing a yellow fluorescent protein is cotransformed with a red fluorescent plasmid, which is either expressing a mutant protein or a CRISPR/Cas9 system for inducing mutations. **B.** Three population mosaic expression analysis. In this genetic interaction study, two different proteins are fused to either sfGFP1-10 (G1) or sfGFP11. Cotransformation of both plasmids yields a subset of bacteria that are green fluorescent. In both schemes, deviation from the fractal pattern of growth would reveal genes involved in colony development and can reveal different patterns of bacterial growth.

## Conclusion

Our observations confirmed that a single bacterium is the foundation of our mixed colony phenotypes. Cotransformation of several plasmids was readily observed with as little as 0.1 ng of each plasmid, a concentration that is remarkably lower than what is commonly used for cloning or retransformation cloned DNA. Thus, many existing plasmid preparations likely contain multiple types of plasmids, some containing mutations ^23, 24^. When retransformed at too high a concentration, resulting bacterial colonies may have preferentially selected mutant plasmids capable of outcompeting the original one. Our lab routinely sequences new subclones from insert ligations with primers that can reveal multiple sequences, i.e. multiple insertion events from distinct clones, which are accidently grown at the same time by the procsss of contransformation or contamination described above. This analysis provides us with the basis for discarding clones that have multiple sequence traces at any position in the insert. In addition, we recognize that bacterial colonies that are not completely circular can arise from two neighboring colonies. To limit DNA from multiple plasmids in a preparation, we select colonies that are completely circular. Finally, we foresee potential of our fluorescent protein expression system to replace the blue/white subcloning techniques with a fluorescent/non-fluorescent cloning methodology. Typical blue/white selection is performed by adding X-gal or IPTG/X-gal to bacterial plates followed by the plating of DNA transformation. X-gal diluted in N, N Dimethylformamide, can reduce the growth rate of bacteria and thus reduce plasmid yield. In contrast, nothing needs to be added to our Hi→Teal, Hi→Venus, Hi→Cherry and Hi→sfGFP vector transformations in order to produce bright fluorescent colonies that with an observable color in white light after an overnight growth (Figure S9). When an insert is cloned into a polylinker disrupting its open reading frame, the fluorescence is abolished yielding white colonies. We believe that the Hi→Cherry fluorescing colonies yielded the most contrast with the white colonies insert containing colonies (Figure S9).

We provide a profound set of data to reveal properties of bacterial transformation that challenge how we perceive molecular DNA cloning and unmask fractal growth patterns of *E. coli* colonies from a single bacterium founder to a whole colony. Importantly, we provide a platform to further pursue either avenue of investigation. In particular, these assays could further the understanding of genes involved in growth and replication patterns of *E. coli* and potentially other bacteria, and may yield clues to the development of microbial biofilms, relevant to human pathogenesis ^25^.

## Acknowledgments

We thank Ivan Rodriguez, Thomas Bozza, Diana Bratu, Carmen Melendez-Vasquez and Anna-Katerina Hadjantonakis for helpful insights throughout the data collection phase. Also, Ivan Diaz, Assistant Professor, Division of Biostatistics Department of Healthcare Policy & Research, Weill Cornell Medical School for helping with the analysis. Many thanks to the members of the lab, especially Eugene Lempert, Ute Schwinghammer, Irena Parvanova, Charlotte D’Hulst for providing thoughtful comments. This work was supported by Research Centers in Minority Institutions Program grant from the National Institute on Minority Health and Health Disparities (MD007599), NIH SC1 GM088114 (P.F), and CTSC UL1 TR000457-06 (I.D.). We would like to dedicate our manuscript to Roger Tsien, who made it possible to peek inside cells with an array of fluorescent technologies.

## Materials and Methods

### DNA transformations

All DNA transformations were carried out with 100 mM CaCl_2_ competent DH5 α*E.coli*. In general: DNA incubations were performed with cells on ice for 30 minutes, followed by a 1 minute heat shock at 42°C, then a 1 minute incubation step back on ice. For carbenicillin (*lam* plasmids) plates (1.5% agar), 600 Ιl of media was added before plating. For kanamycin (*kan* plasmids) plates (1.5% agar), 600 Ιl of media was added, followed by 30 minute incubation at 37C prior to plating. For pGLO induction with 2% L-arabinose was performed after overnight growth for 6 hrs (http://www.bio-rad.com/en-us/product/pglo-bacterial-transformation-kitInduction) All images (except movies) were taken with a 2.5x, 5x, 10x or 20X/0.5 Plan-Neofluar Zeiss lens using an LSM510 confocal microscope.

### Cotransformation of 10 Venus plasmids

Ten Venus plasmids (V1-V10) plus one Cherry plasmid (D355-2), each at 1 ng/μl (∼11 ng total) plus 50 1l CaCl_2_, ice 25 minutes, heat shock 1 minute 15 seconds, ice 1 minute. 1 ng of Teal plasmid was then added (D378-1), then addition of 600 1l of media and plating. 12 colonies were picked (10 with observable Venus fluorescence and 2 without). Only a portion of each colony assayed by PCR was picked.

### Movies

A mixed colony was picked into a well of a 96-well plate and grown overnight in 100 1l of 2XYT media. 1 1l was taken out and diluted into 1 ml of 2xYT media. 1 1l was again taken out of this dilution and and diluted into 1 ml of 2xYT media. 300 1l of it was dispensed on a agarose plate. The next day, a tip was dipped in the colony, then swirled in 1ml of 2xYT media. 10 1l -20 1l of this media was placed in the center of a small agarose plate. Small plates were made of a 4.5% agarose main bottom layer topped with a superficial layer of 3% agarose. The plates were placed on a heated microscope stage at 33-35C and cover-slipped. Images were taken with 20X/0.5 Plan-Neofluar Zeiss lens on a LSM 510 Zeiss Confocal Microscope at 4x optical zoom. The stage was moved up in the Z position by about 20 uM every 20 minutes to manually focus.

## Supplementary Figure Legends

**Fig. S1.**
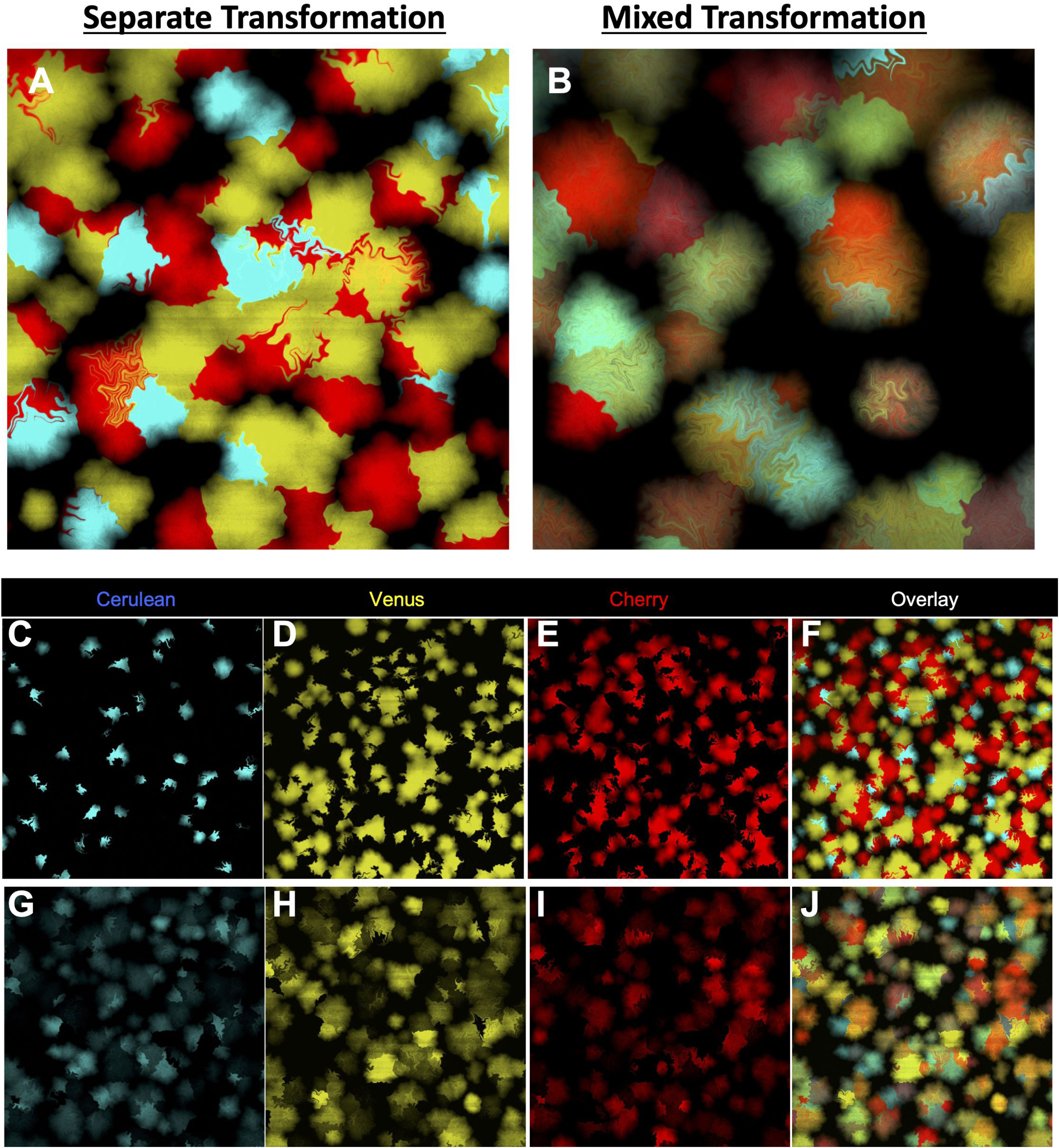
Mixed colonies are not separable into components upon replating. **A, C-F**, Replating of mixture of Cerulean, Venus and Cherry fluorescing colonies at high density still show separation of fluorescence. **A and F** are overlay. **B, G-J**, by contrast, replating of a single mixed Cerulean, Venus and mCherry expressing colony at high density retains the fluorescent coexpression in all resulting colonies. **B and J** are overlay.

**Fig. S2.**
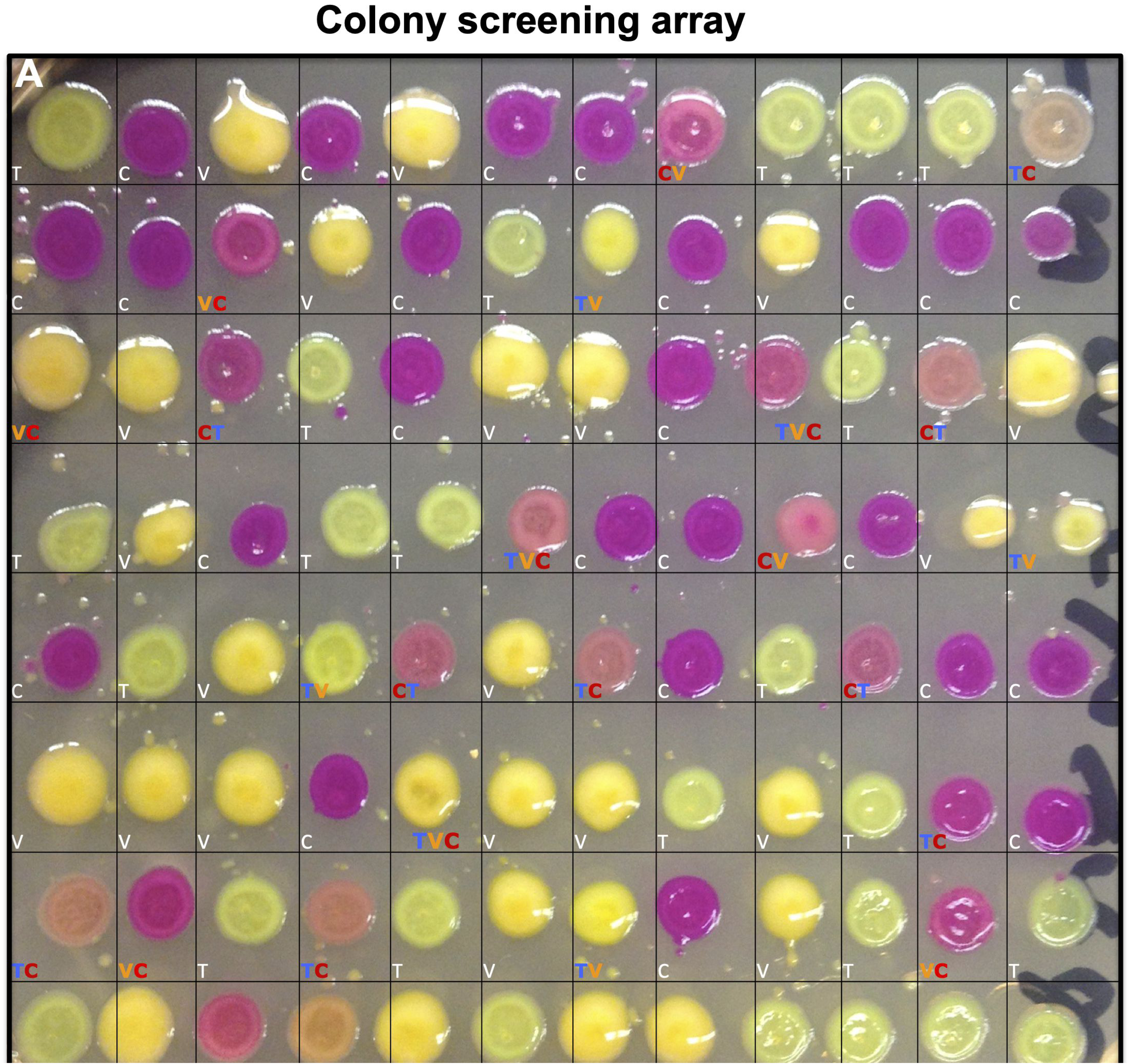
Individual bacterial colonies arrayed for screening of fluorescent coexpression. **A.** Single colonies were picked at random after transformations and grown overnight in a 96 well plate with 100 μl of selective media. The following day, 1 μl from each well are arrayed on a bacterial plate and grown for 12 hrs at 37 C followed by growth on the bench for 1-2 days. Hi→Teal (T), Hi→Venus (V), and Hi→Cherry (C) bacterial growths are readily visible. Double and triple coexpressing bacterial growths are also visible (TV, VC, CT, and TVC) and were assigned by confocal microscopy. For the double transformants we ranked the higher fluorescent output first in the annotations (see Fig. S3 and Data File S1).

**Fig. S3.**
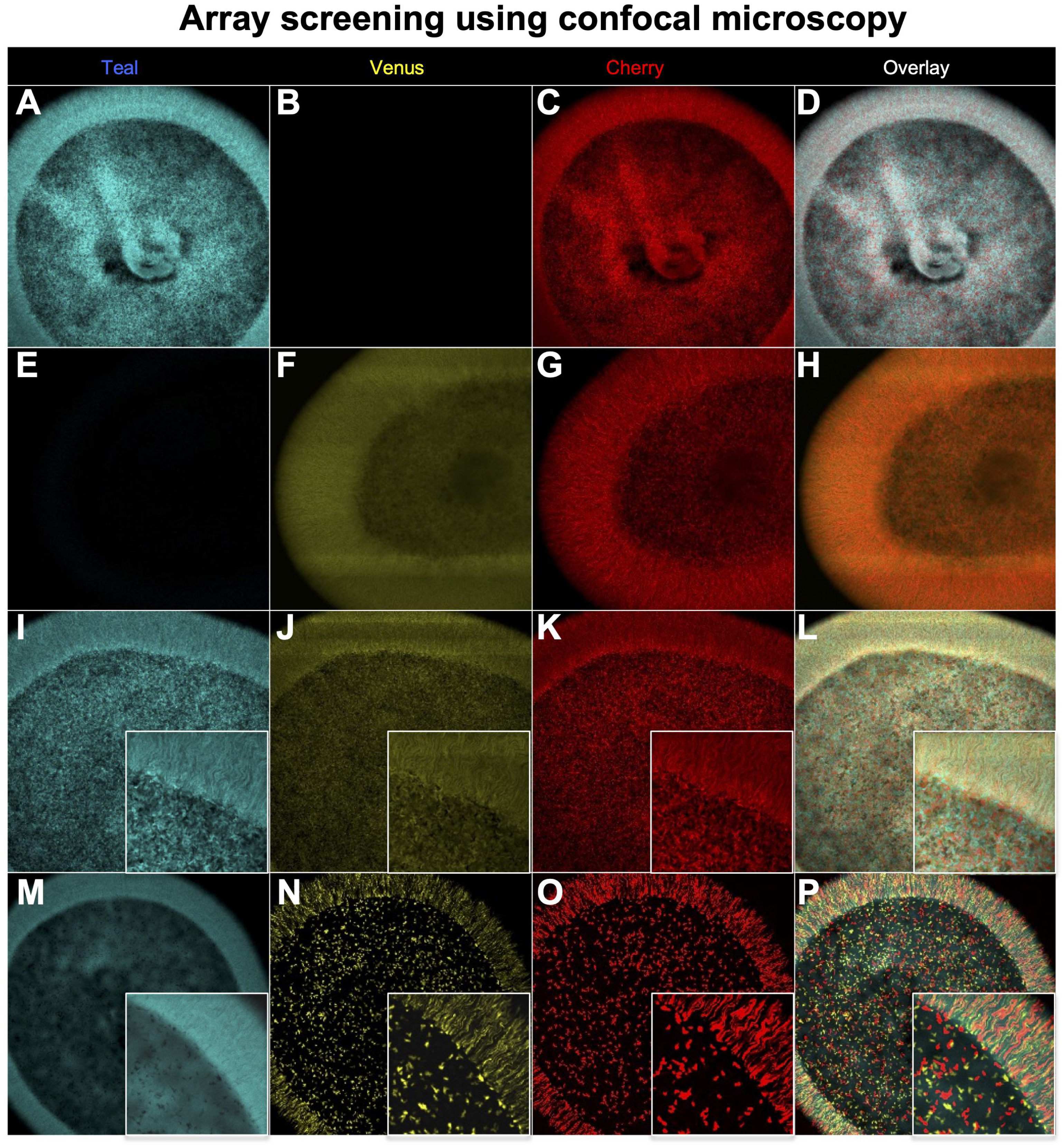
Screening of bacterial growths for coexpression by fluorescent microscopy. **A-C,** A Hi→Teal (T) and Hi→Cherry (C) coexpressing bacterial growth (TC). **D** is overlay showing complete overlap between Teal and Venus fluorescence. **E-G,** A Hi→Venus (V) and Hi→Cherry (C) coexpressing bacterial growth (VC). **H** is overlay showing complete overlap between Venus and Cherry fluorescence. **I-K,** A Hi→Teal, Hi→Venus, and Hi→Cherry coexpressing bacterial growth (TVC). **L** is overlay showing complete overlap between Teal, Venus and Cherry fluorescence. **M-O,** A Hi→Teal bacterial growth contaminated with Hi→Venus and Hi→Cherry bacteria. **P** is overlay showing contamination having different morphology than the main bacterial content. High magnification within **M-P** reveals the details of bacterial contamination: Hi→Venus and Hi→Cherry appear as holes in Teal fluorescent image and streaks at the border of the bacterial growth.

**Fig. S4.**
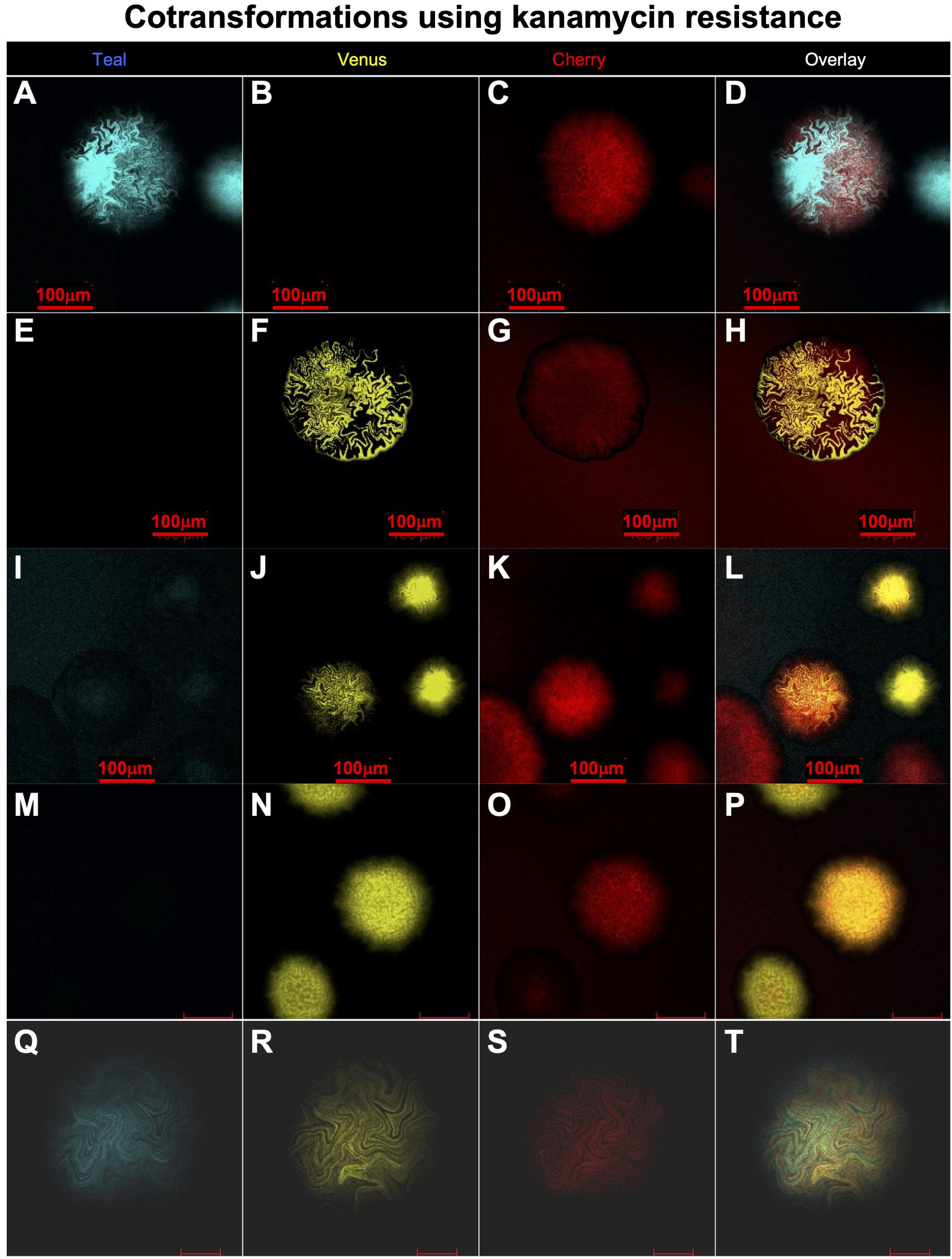
Cotransformation of plasmids with two different resistances. **A-C,** A Kanamycin selected single colony from transformation with Hi→Teal (*lam*) and Lo→Cherry (*kan*). **D** is overlay mosaicism of Teal fluorescence. **E-G,** A kanamycin selected single colony from transformation with Hi→Venus (*lam*) and Lo→Cherry (*kan*). **H** is overlay showing mosaicism of Venus fluorescence. **I-K,** Kanamycin selected single colonies from transformation with Hi→Venus (*lam*) and Lo→Cherry (*kan*). **J** shows mosaicism of unselected *(lam)* Venus fluorescence in one colony as well as high expression in several other colonies whereas **K** shows selected *(kan)* Cherry fluorescence in all colonies. **L** is overlay. **M-O,** Carbenicilin selected colonies from a Hi→Venus (*lam*) and Lo→Cherry (*kan*) positive colony. **N** shows all selected *(lam)* colonies with Venus fluorescence and **O** shows mosaic unselected *(kan)* Cherry expression in one colony. **P** is overlay. **Q-S,** A kanamycin selected colony cotransformed with 4 plasmids: Hi→Teal (*lam*), Hi→Venus (*lam*), Hi→Cherry (*lam*) and Lo→Cherry (*kan*). Fluorescence in **S** is from Hi→Cherry (*lam*) and not from Lo→Cherry (*kan*) for two reasons, first the laser excitation intensity was reduced to a level that only detects Hi→Cherry and not Lo→Cherry and second the pattern of mosaicism is consistent with what is observed when three plasmids expressing three different fluorescent proteins under the same selective resistance. **T** is overlay.

**Fig. S5.**
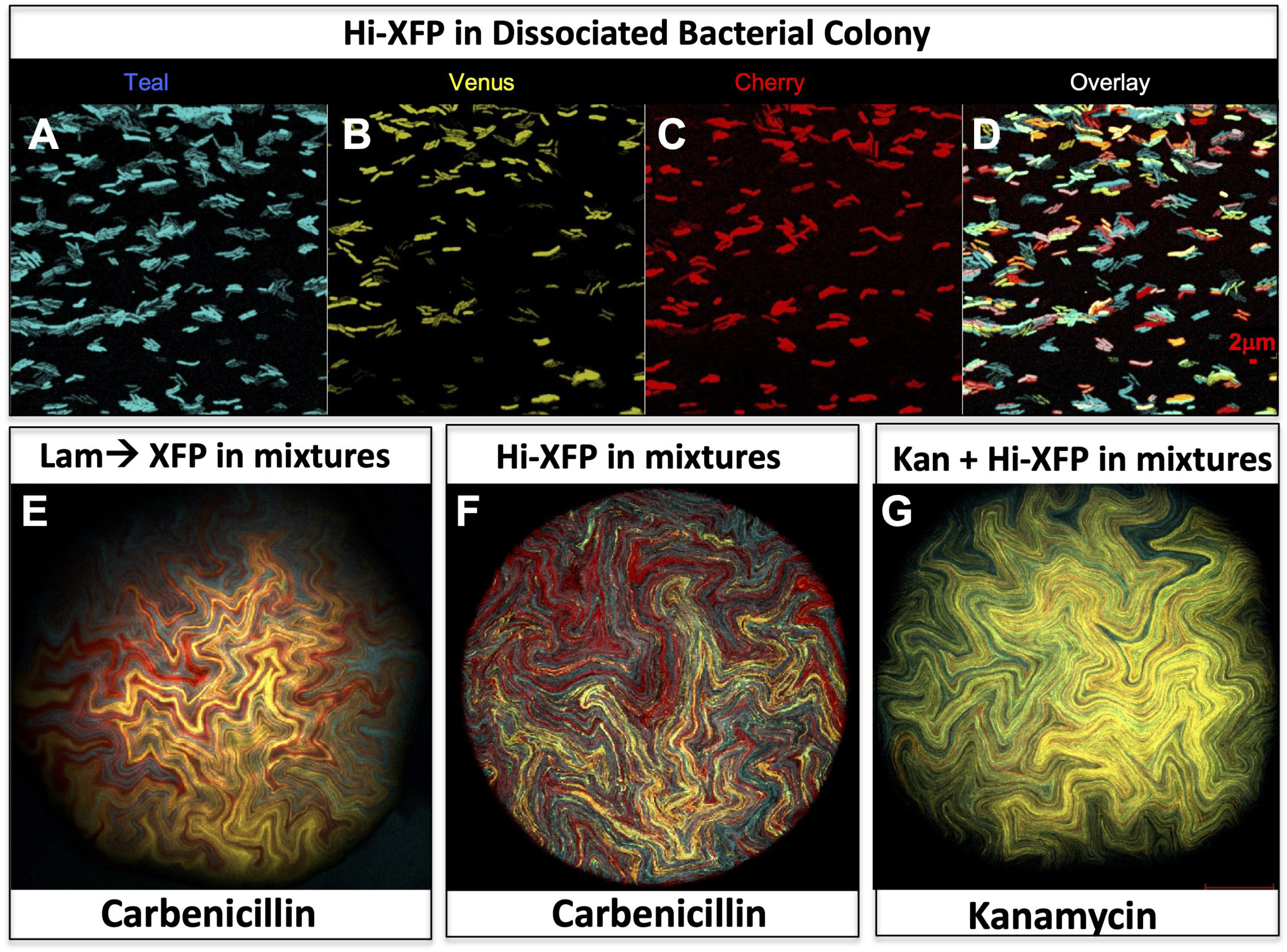
Cotransformation of multiple fluorescent plasmids with or without selection have similar mosaicism. **A-D.** triple fluorescent bacterium from a triple fluorescent colony expressing Hi→Teal, Hi→Venus, and Hi→Cherry and Overlay. Scale bar at 2μm**. E.** A carbenicillin resistant triple fluorescent colony (Lam→Teal, Lam→Venus, and Lam→Cherry) shows mosaic fluorescence (Projected Z-Stack). **F.** A carbenicillin resistant triple fluorescent colony (Hi→Teal, Hi→Venus, and Hi→Cherry) shows mosaic fluorescence (Projected Z-Stack) with single bacterium resolution. **F**. A kanamycin resistant triple fluorescent colony (Hi→Teal, Hi→Venus, and Hi→Cherry: not Carb selected) shows mosaic fluorescence (Projected Z-Stack) without single bacterium resolution; Same colony as in **Fig. S4.** panel **T**. Despite kanamycin resistance from a Lo→Cherry (*kan*) plasmid, still reveals strong fluorescence similar to colony in **A. E-F,** 500μm bacterial colony.

**Fig. S6.**
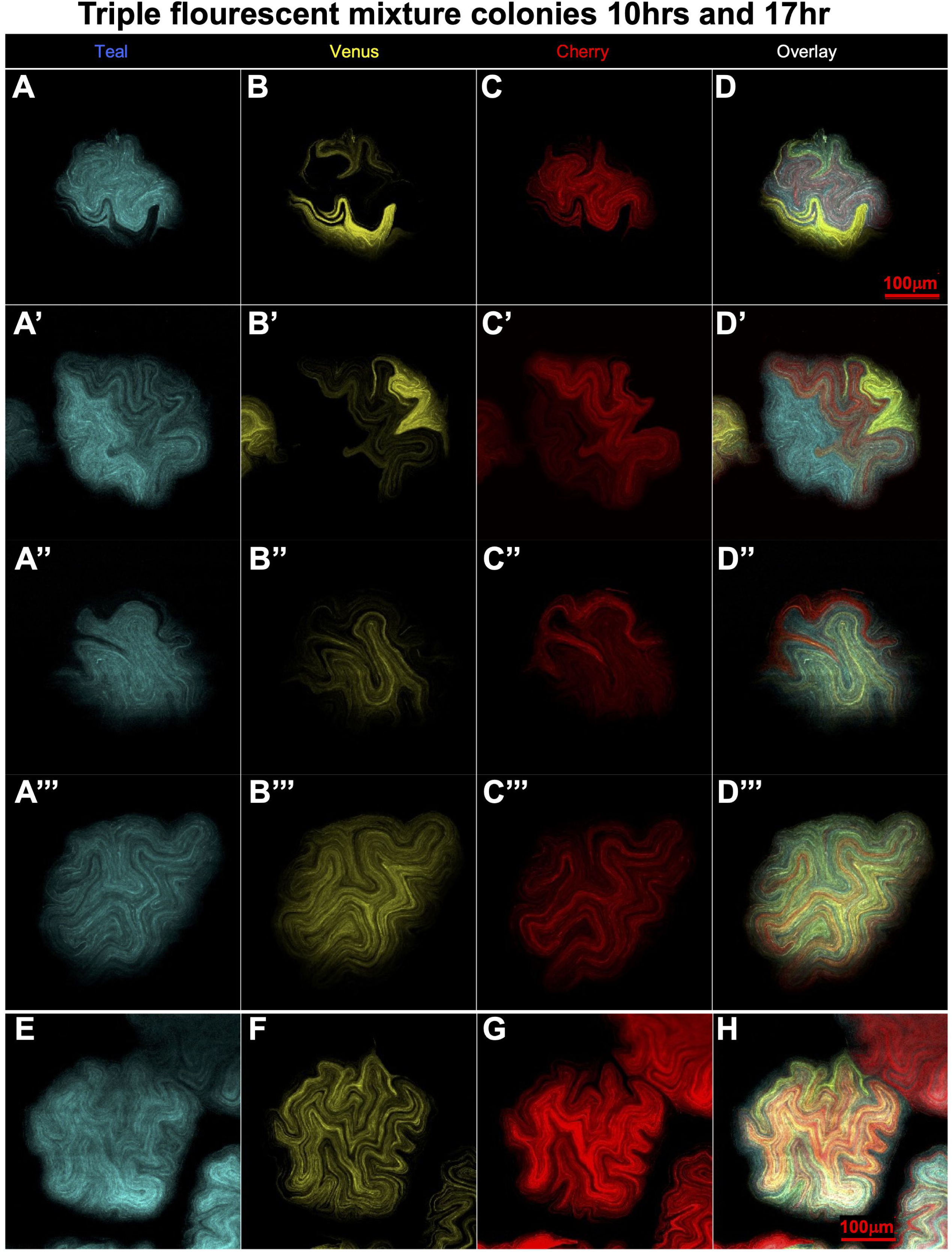
Mosaic fluorescence observed at 10hr and 17hr of growth. **A-D, A’-D’, A’’-D’’, and A’’’-D’’’** from 4 additional triple fluorescent expressing Hi→Teal, Hi→Venus, and Hi→Cherry at 10 hrs of growth and 150 µm in width. **D-D’’’** are overlays. **E-G** is from a triple fluorescent colony expressing Hi→Teal, Hi→Venus, and Hi→Cherry at 17 hrs of growth and 200 µm in width. **H** is overlay. All five colonies reveal same mosaicism as found in older colonies.

**Fig. S7.**
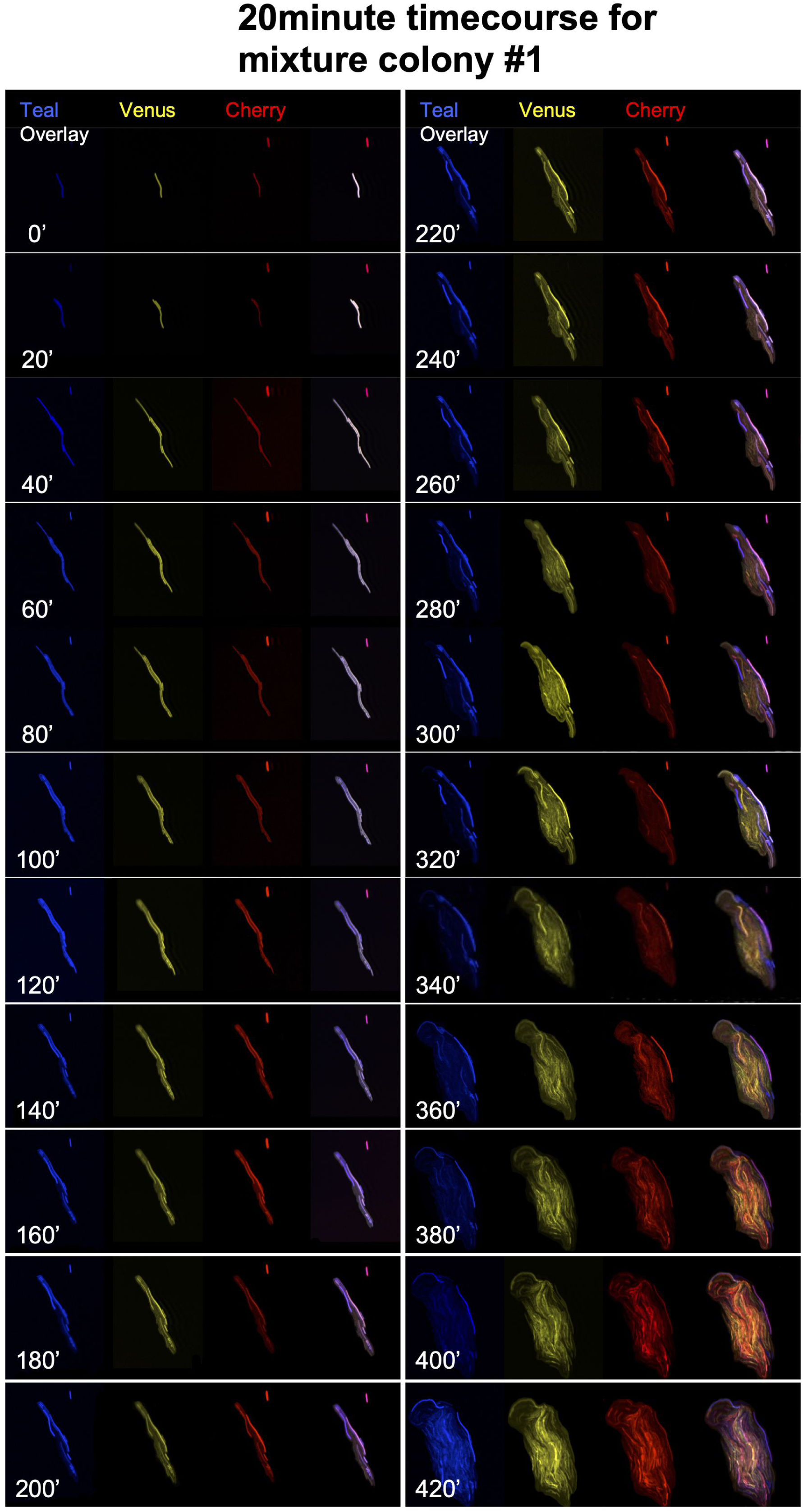
Time course of triple fluorescent colony development 0 minutes to 420 minutes. A triple fluorescent bacterium (A) expressing Hi→Teal, Hi→Venus, and Hi→Cherry was imaged every 20 minutes for 420 minutes. See also **Movie S1A and S1B.** Mosaicism is quickly revealed after a few cell divisions.

**Fig. S8.**
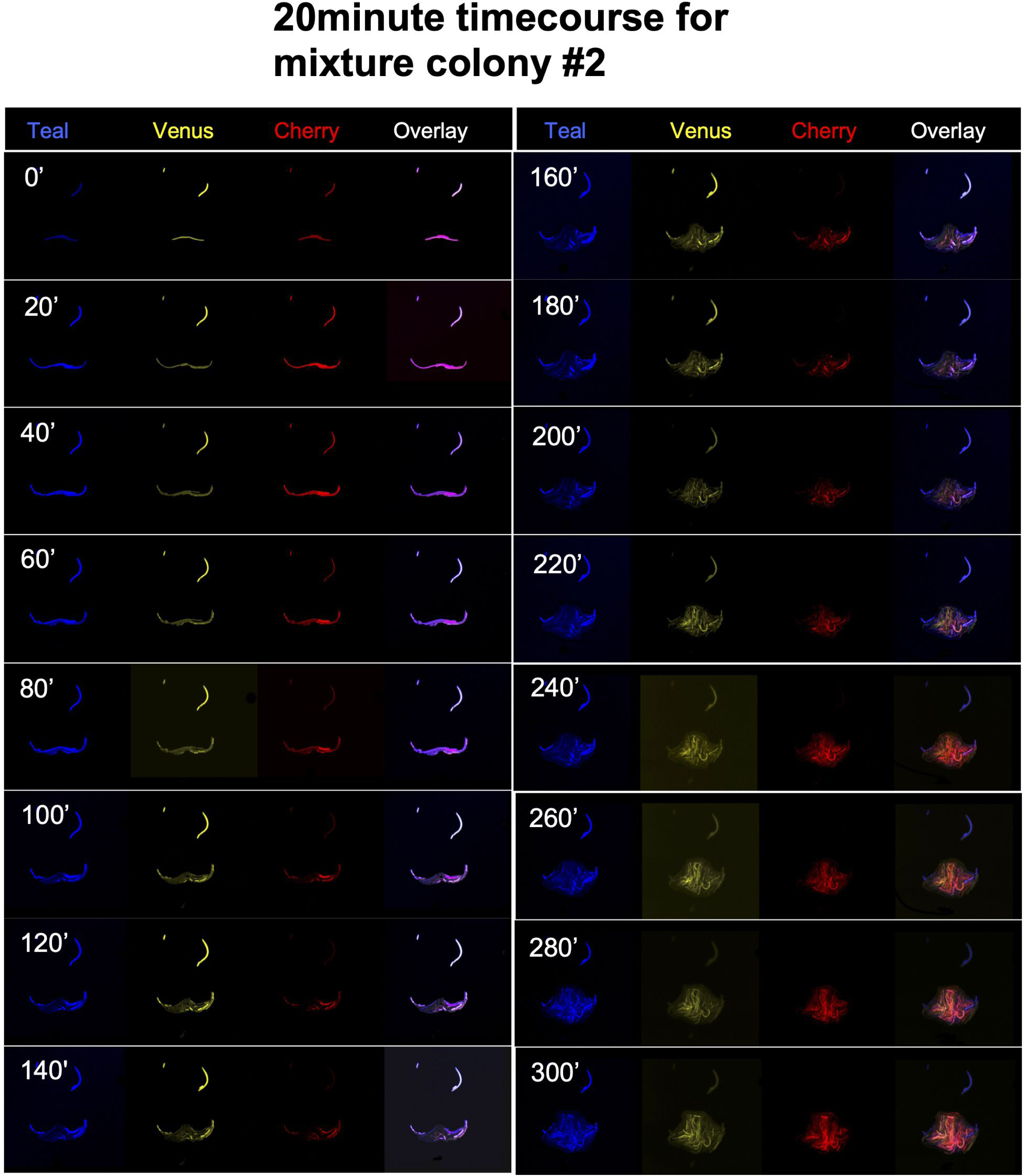
Time course of triple fluorescent colony development 0 minutes to 300 minutes. A second triple fluorescent bacterium (B) expressing Hi→Teal, Hi→Venus, and Hi→Cherry was imaged every 20 minutes for 300 minutes. See also **Movie S2.** Mosaicism is quickly revealed after a few cell divisions.

**Fig. S9.**
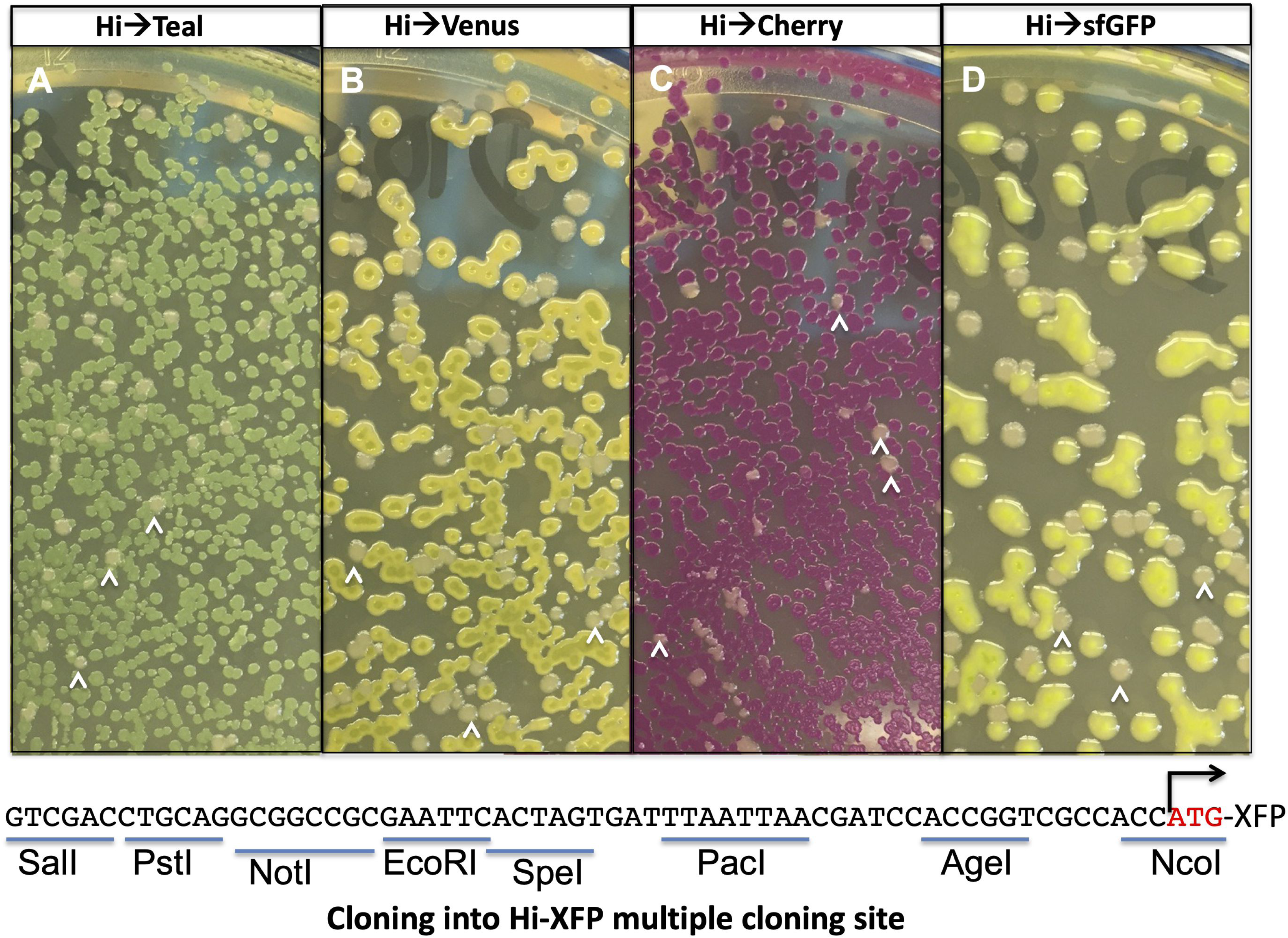
Colorimetric assay for cloning insert DNA. B) Hi→Teal, B) Hi→Venus, C) Hi→Cherry and D) Hi→sfGFP colonies observed in white light after overnight growth on agar plates. If a 1kb SpeI DNA fragment (an insert) is cloned into any of these vectors at the SpeI site in the common polylinker, then white colonies emerge on the plate (white arrowheads). PCR screening of white colonies confirms that they indeed carry the insert. Thus, a colorimetric assay can be used without fluorescent light to identify subcloned DNA fragments.

**Movie. S1A, S1B. Triple fluorescent bacterium. (S1A). Teal FP (blue), Venus FP (yellow), Cherry FP (red) and Overlay views for a s**ingle bacterial rod that grew over time and were typically longer than a typical 1x 2 µm rod. Note that a Hi→Teal and Hi→Cherry coexpressing bacterium in the same field of you did not replicate and would have been engulfed by the growing colony had it been allowed to grow longer. It is not clear if this non-replicating bacterium is dead or unable to replicate on the plate. These “contaminating” bacteria may be what are represented in panel **Fig. S3. P. (S1B) Overlay only for split image in (S1).**

**Movie. S2. Triple fluorescent bacterium (B) Teal FP (blue), Venus FP (yellow), Cherry FP (red) and Overlay views for a s**ingle bacterial rod that grew over time and were typically longer than a typical 1x 2µm rod. Note that two other triple fluorescent bacterium did not replicate would have been engulfed by the growing colony. It is not clear if these non-replicating bacteria are dead or unable to replicate on the plate. These “contaminating” bacteria may be what are represented in panel **Fig. S3. P.**

**Movie. S3.** Z-stack projection of a triple fluorescent colony (Hi→Teal, Hi→Venus, and Hi→Cherry) reveals fluorescence in individual colonies in several planes.

**Movie. S4.** Z-stack projection of a second triple fluorescent colony (Hi→Teal, Hi→Venus, and Hi→Cherry) reveals fluorescence in individual colonies in several planes.

## Table Legends

**Table S1.**
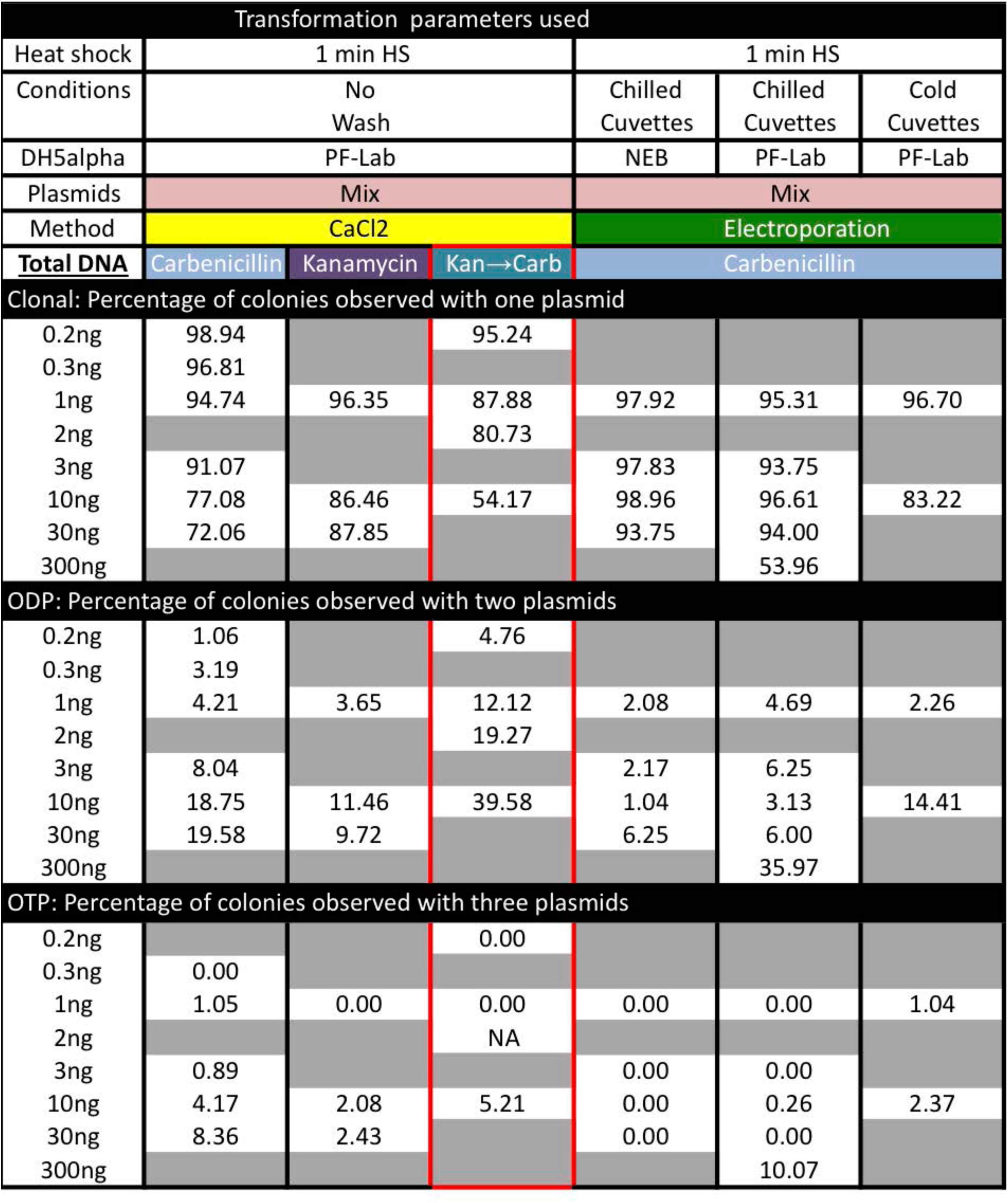
Summary of cotransformation rates with multiple plasmids under different parameters. The Kan→Carb column reflects concentrations for Carb only. In addition to this, 1 ng of Kan plasmid was added during cotransformation experiments.

Transformation parameters used
| Heat shock | 1 min HS |  |  | 1 min HS |  |  |
| --- | --- | --- | --- | --- | --- | --- |
| Conditions | No Wash |  |  | Chilled Cuvettes | Chilled Cuvettes | Cold Cuvettes |
| DH5alpha | PF-Lab |  |  | NEB | PF-Lab | PF-Lab |
| Plasmids | Mix |  |  | Mix |  |  |
| Method | CaCl2 |  |  | Electroporation |  |  |
| Total DNA | Carbenicillin | Kanamycin | Kan→Carb | Carbenicillin |  |  |
| Clonal: Percentage of colonies observed with one plasmid |  |  |  |  |  |  |
| 0.2ng | 98.94 |  | 95.24 |  |  |  |
| 0.3ng | 96.81 |  |  |  |  |  |
| 1ng | 94.74 | 96.35 | 87.88 | 97.92 | 95.31 | 96.70 |
| 2ng |  |  | 80.73 |  |  |  |
| 3ng | 91.07 |  |  | 97.83 | 93.75 |  |
| 10ng | 77.08 | 86.46 | 54.17 | 98.96 | 96.61 | 83.22 |
| 30ng | 72.06 | 87.85 |  | 93.75 | 94.00 |  |
| 300ng |  |  |  |  | 53.96 |  |
| ODP: Percentage of colonies observed with two plasmids |  |  |  |  |  |  |
| 0.2ng | 1.06 |  | 4.76 |  |  |  |
| 0.3ng | 3.19 |  |  |  |  |  |
| 1ng | 4.21 | 3.65 | 12.12 | 2.08 | 4.69 | 2.26 |
| 2ng |  |  | 19.27 |  |  |  |
| 3ng | 8.04 |  |  | 2.17 | 6.25 |  |
| 10ng | 18.75 | 11.46 | 39.58 | 1.04 | 3.13 | 14.41 |
| 30ng | 19.58 | 9.72 |  | 6.25 | 6.00 |  |
| 300ng |  |  |  |  | 35.97 |  |
| OTP: Percentage of colonies observed with three plasmids |  |  |  |  |  |  |
| 0.2ng |  |  | 0.00 |  |  |  |
| 0.3ng | 0.00 |  |  |  |  |  |
| 1ng | 1.05 | 0.00 | 0.00 | 0.00 | 0.00 | 1.04 |
| 2ng |  |  | NA |  |  |  |
| 3ng | 0.89 |  |  | 0.00 | 0.00 |  |
| 10ng | 4.17 | 2.08 | 5.21 | 0.00 | 0.26 | 2.37 |
| 30ng | 8.36 | 2.43 |  | 0.00 | 0.00 |  |
| 300ng |  |  |  |  | 10.07 |  |

**Table S2.** Summary of cotransformation rates with multiple plasmids with wash step added after heat shock.

| Transformation parameters used |  |  |  |  |  |  |
| --- | --- | --- | --- | --- | --- | --- |
| Heat shock | 1 min HS | 1 min HS | 1 min HS | 1 min HS | Short HS | Short HS |
| Conditions | Cold | Cold | RT | RT | RT | RT |
| Wash | CaCl2 | CaCl2 | Water | Water | Water | Water |
| DH5alpha | PF-Lab |  |  |  |  |  |
| Plasmids | Mix | Sep | Mix | Sep | Mix | Sep |
| Method | CaCl2 |  |  |  |  |  |
| **<u>Total DNA</u>** | Carbenicillin |  |  |  |  |  |
| Clonal: Percentage of colonies observed with one plasmid |  |  |  |  |  |  |
| 30ng | 72.92 | 98.96 | 89.58 | 100.00 | 59.52 | 96.87 |
| ODP: Percentage of colonies observed with two plasmids |  |  |  |  |  |  |
| 30ng | 26.04 | 1.04 | 9.38 | 0.00 | 32.14 | 3.13 |
| OTP: Percentage of colonies observed with three plasmids |  |  |  |  |  |  |
| 30ng | 1.04 | 0.00 | 1.04 | 0.00 | 8.33 | 0.00 |

## Supplemental information inventory

### Supplemental Figures and Legends

S1 (related to Figure 1 and 2): Three colony mixture versus a triple fluorescent colony.

S2 (related to Table 1): Colony screening array.

S3 (related to Table 1): Screening colony array using confocal microscopy.

S4 (related to Table 1): Cotransformations using kanamycin resistance.

S5 (related to Figure 5 and Table 1): Triple fluorescent mixture colonies.

S6 (related to Figure 6): Triple fluorescent mixture colonies: 10hr & 17hr.

S7 (related to Figure 7): 20 minute time course for mixture colony #1.

S8 (related to Figure 7): 20 minute time course for mixture colony #2.

S9 (related to Figure 4): Colormetric screening assay for inserts

### Supplemental Table

S1 (related to Table 1): Calculated cotransformation rates with probabilities matrix.

S2 (related to Table 1): Cotransformation rates under new conditions.

### Supplemental Movies

S1A (related to Figure 7): Split fluorescence movie of time course for mixture colony #1.

S1B (related to Figure 7): fluorescence movie of time course for mixture colony #1.

S2 (related to Figure 7 and S7): Split fluorescence movie of time course for mixture colony #2.

S3 (related to Figure 5 and S5): Z-stack projection of a triple fluorescent colony.

S4 (related to Figure 5 and S5): Z-stack projection of a second triple fluorescent colony.

### Supplemental Data File

S1 (related to Figure 4, 5, 6, 7, S2, S3, S4 and Table S1): Raw data for cotransformation analysis.

S2 (related to Table S1): Calculated probabilities matrix.

## Data File S2

We have the following distribution of colors for plasmids C (Cherry), T (Teal) and V (Venus):

| # of types | # of colors observed |  |  | Calculated probabilities |
| --- | --- | --- | --- | --- |
|  | 1 | 2 | 3 |  |
| 1 plasmid | **w11** | 0 | 0 | **Q1** |
| 2 plasmids | w21 | **w22** | 0 | **Q2** |
| 3 plasmids | w31 | w32 | **w33** | **Q3** |
| Observed Probabilities | **P1** | **P2** | **P3** | 1 |

**For the values of w11, w21, w22, w31, w32 and w33 the conditional probabilities were surmised as follows:**

1) There are three types of w11: C, T and V

2) Given a C, T or V event there is an equal chance that a w21 event or w22 event has occurred:

There are three types of w21 events: CC, TT and VV
There are three types of w22 events: CV, CT, and TV

3a) Given there are CC, TT or VV events then,

- there are three types of w31 events: CCC, TTT and VVV
- there are six types of w32 events: CCT, CCV, TTV, TTC, VVC and VVT
- there are zero w33 events
3b) Given there are VT, VC or CT events then,
- there are zero w31 events
- there are six types of w32 events: VTT, VVT, VCC, VVC, CTT, and CCT
- there are three types of w33 events VTC, VCT or CTV.
Thus there are just as many w33 events as w31 and 4 fold more w32 events than w31 or w33 events.

**The equations to compute theoretical single, double and triple positives:**

P1 is equal to our observed single positives (putatively clonal), w11
P2 is equal to our observed double positives, w22
P3 is equal to our observed triple positives*#, w33
Q1 will be the estimated single positives (clonal)
Q2 will be the estimated double positives
Q3 will be the estimated triple positives

Therefore:

Q3= w33 (w31) + 4x w33 (w32) + w33 (P3)= 6x P3
**Q3= 6 x P3**

Q2= w21 (w22) + w22 (P2) = 2 x P2, but we must subtract triple events that look like double events (w32) or ?4x P3
**Q2=2x P2C4x P3**

Q1= w11 (P1), but we must subtract double events that look like single events, w21 (w22=P2), and triple events that look like single events, w31 (w33= P3)
**Q1= P1CP2CP3**

**Q1+ Q2 +Q3=P1 + (2xP2CP2)+ (6x P3C4x P3CP3) = P1 +P2 + P3**

**\*Based on the fold reduction in clonality once a w33 event is observed (high number of unobserved triple events), we surmise that 4 plasmid events: w41, 42, w43, begin to occur concomitantly. Thus w33 events signal the beginning of a precipitous drop in clonality.**

**#For Kan◊Carb experiments there was a single four plasmid event. This event was not added to P3 in Table 1 or Table S1. If this event is added to the P3 percentage, then Q3=37.49%, Q2=54.17% and Q1=8.34%**

